# A Modular Machine Learning Framework for Small-Molecule Drug Repurposing Based on Organ Permeability, Target Binding, and Biomarker Modulation

**DOI:** 10.64898/2026.04.14.718573

**Authors:** Harkirat Singh Arora, Saroj Dhakal, Yorgos Psarellis, Sriram Chandrasekaran, Panteleimon D. Mavroudis, Nikhil Pillai

## Abstract

With nearly 90% of drug candidates failing in clinical trials due to poor efficacy or toxicity, drug repurposing has emerged as a vital strategy to accelerate the delivery of life-saving treatments. However, most current drug repurposing approaches fail to account for the physiological barriers and downstream biological impacts that dictate therapeutic success. To bridge this gap, we present SCOUT (Screening Candidates via Organ Uptake and Target-binding), a modular machine learning-driven framework for drug repurposing by simultaneously modeling organ permeability, drug-target binding, and biomarker modulation. Unlike conventional repurposing efforts that rely on single-point predictions or disconnected steps, SCOUT integrates these factors into a unified predictive funnel, significantly reducing trial-and-error and prioritizing candidates with the highest probability of therapeutic relevance. As a proof of concept, we applied SCOUT to Alzheimer’s disease, where blood-brain barrier (BBB) penetration is an important hurdle. The framework first predicts the unbound brain-to-plasma partition coefficient, Kpuu, using SMILES-based embeddings and achieved robust performance (Accuracy: 0.90, Recall: 0.94), then integrates machine learning to predict binding affinity with BACE1, a key Alzheimer’s target that was used in this work as a proof of concept. By combining these modules, SCOUT reduced the candidate search space by 99.9% and identified hits with diverse mechanisms of action. Critically, SCOUT extends beyond hit identification by employing mechanistic modeling to simulate how these candidates might modulate specific disease biomarkers (e.g. Amyloid-beta (Aβ) peptides). This hybrid approach ensures repurposed candidates are not only chemically viable but also physiologically active, providing a rational and resource-efficient pipeline for drug development.

## Introduction

Drug discovery is a prohibitively expensive and time-consuming endeavor, typically requiring over a decade and an investment exceeding $2–3 billion per successful agent [1]. Despite the significant investment, nearly 90% of candidates fail in clinical trials, often due to insufficient efficacy or unmanageable toxicity [2]. Drug repurposing is the process of identifying new therapeutic uses for approved or investigational drugs and provides an attractive and economically viable strategy [3]. By leveraging existing safety data, repurposed drugs could possibly bypass preclinical development and Phase I trials and enter directly into Phase II clinical testing to confirm therapeutic efficacy [4]–[7]. Consequently, nearly 30% of current FDA-approved therapies are the result of repositioning, underscoring the immense potential of this strategy [8], [9].

To capitalize on this potential, researchers have increasingly turned to computational methods to scan the vast chemical and biological space for viable candidates. Early efforts focused on network-based approaches, utilizing annotations of drug such as chemical structure, known side effects, etc., and their corresponding MeSH (Medical Subject Headings) terms, drug-drug and disease-disease similarity, drug-disease associations, gene-expression profiles, and proteomic proximity to identify overlap between drug targets and disease pathways [10]–[14]. More recently, the advent of machine learning (ML) has enabled more sophisticated data-driven predictive modeling. For instance, tools like DRIAD have quantified molecular associations in Alzheimer’s pathology [15], while frameworks such as DrugRepo have integrated similarity scores to screen drugs across hundreds of diseases [16]. Other frameworks, like SperoPredictor, have combined ML with molecular docking to rapidly identify therapeutics for emergent crises like COVID-19 [17].

For our proof-of-concept, we apply the SCOUT (Screening Candidates via Organ Uptake and Target-binding) framework to Alzheimer’s disease, where many times the blood-brain barrier (BBB) serves as an important factor governing therapeutic success. Traditionally, computational models have relied on logBB (the log ratio of total drug concentration in the brain vs. plasma) to quantify permeability. High-performing models using Random Forests [18] and deep learning architectures like MegaMolBART [19] have achieved impressive AUC scores of 0.95 and 0.88 respectively for logBB prediction. However, logBB is increasingly viewed as an incomplete metric because it does not distinguish between protein-bound and pharmacologically active drug concentrations. Consequently, as per the free-drug hypothesis [20], the unbound brain-to-plasma partition coefficient (Kp,uu) has emerged as the gold standard for predicting true penetration [21]–[23]. Despite its importance, predicting Kp,uu remains challenging due to the scarcity of high-quality datasets. Recent efforts have utilized molecular descriptors [24], [25] and solvation energy data [26] to predict Kp,uu, yet these models often struggle with moderate R^2^ values (0.53 - 0.61) or limited categorical accuracy (0.79 - 0.85). SCOUT seeks to advance this field by implementing a high-performance Kp,uu module that serves as the first critical filter in our repurposing funnel.

Small molecule drugs exert their therapeutic influence via perturbation of protein networks. Therefore, modeling drug-protein interactions is critical as it helps in identifying candidate drugs that can bind with a disease-relevant target. In the context of Alzheimer’s disease, BACE1 (Beta-secretase 1) has been extensively investigated in literature for its potential role in the progression of disease via amyloidogenic pathway [27]–[30]. Traditionally, drug-target interactions (DTI) have been modeled using molecular docking, which simulates the physical binding of a drug to a protein’s 3D structure [31]. While accurate, docking is inherently limited by high computational costs and poor scalability, often requiring minutes or hours to process a single interaction. To bypass this bottleneck, in recent years, a substantial progress has been made in using machine learning to predict drug-target interactions at scale [32], [33]. Frameworks like DeepPurpose utilize simplified drug representations (SMILES) and protein sequences to predict drug-protein interactions [34]. More recently, large language models (LLMs) such as Boltz-2 have enhanced the predictive precision by integrating complex 3D structural data [35]. Within the SCOUT framework, we leverage a high-throughput ML approach for DTI prediction, allowing us to rapidly filter thousands of candidates and identify those with the highest probability of BACE1 inhibition, thereby bridging the gap between computational speed and biochemical relevance.

While organ permeability and high-affinity drug-target binding are essential prerequisites for therapeutic action, they do not inherently guarantee a meaningful biological response. The complexity of disease pathophysiology often involves compensatory feedback loops where inhibiting a single target may yield insufficient changes in downstream signaling. Crucially, the pharmacokinetic (PK) profile i.e. the magnitude and duration of drug exposure at the site of action serves as the primary determinant of a drug’s sustained biological effect. Mechanistic modeling addresses this critical gap by integrating systems biology with mathematical frameworks, allowing us to simulate the dynamic interplay between drug exposure and the modulation of disease-specific biomarkers. In the context of Alzheimer’s disease, Quantitative Systems Pharmacology (QSP) models have become instrumental in predicting therapeutic outcomes for antibodies, BACE1 inhibitors, and γ-secretase inhibitors [36]. Recent models have specifically simulated the impact of high-profile treatments such as Lecanemab and Aducanumab [37], [38], the latter of which was discontinued in 2024 due to limited clinical benefits, further highlighting the need for models that can predict functional efficacy earlier in the pipeline. Other efforts, such as those by van Maanen et al. [39], have successfully used systems pharmacology to map changes in the amyloid precursor protein (APP) pathway following BACE1 inhibition. Within the SCOUT framework, we bridge the gap between machine learning and physiology by feeding our top candidate hits into these mechanistic models, providing a mechanistic simulation of how repurposed candidates modulate disease biomarker levels (Amyloid-beta (Aβ) peptides).

Despite significant advancements in computational drug repurposing, existing methodologies suffer from three critical bottlenecks. First, most approaches are fragmented, prioritizing a single feature, such as drug-target affinity or drug-drug similarity or drug-disease associations, while ignoring the physiological barriers, like organ permeability, that governs whether a drug reaches its destination. Second, these models often function as black boxes, lacking the mechanistic interpretability required to explain why a specific candidate is viable. Finally, even integrative frameworks (e.g., DrugRepo) fail to bridge the gap between chemical binding and downstream biological impact, leaving the actual modulation of disease biomarkers unaddressed. To overcome these challenges, we introduce SCOUT, a modular, machine learning-driven hybrid framework that unifies organ permeability, drug-target binding, and mechanistic modulation of biomarkers into a single multi-stage predictive funnel. SCOUT ensures that candidates are not only chemically compatible but physiologically relevant. We demonstrated the framework’s power through a proof-of-concept study on Alzheimer’s disease, where SCOUT successfully reduced the candidate search space by 99.9% while identifying hits with diverse mechanisms of action. Further, SCOUT provides interpretable insights into model predictions of why certain candidates have higher probability of repurposing success than others. By coupling machine learning with mechanistic modeling, we provide a rational, interpretable, and highly resource-efficient pathway for discovering the next generation of therapeutics.

## Results

### Integration of Machine Learning and Mechanistic Modeling to develop Hybrid and Modular Framework for Drug Repurposing

Our proposed framework, SCOUT is built upon the integration of two distinct computational arms: a Machine Learning (ML) screening arm for high-throughput candidate identification from a small molecule drug library and a Mechanistic Modeling arm for functional validation of these hits via simulating modulation of downstream biomarker levels. The schematic of the proposed framework is shown in Figure 1.

**Figure 1.**
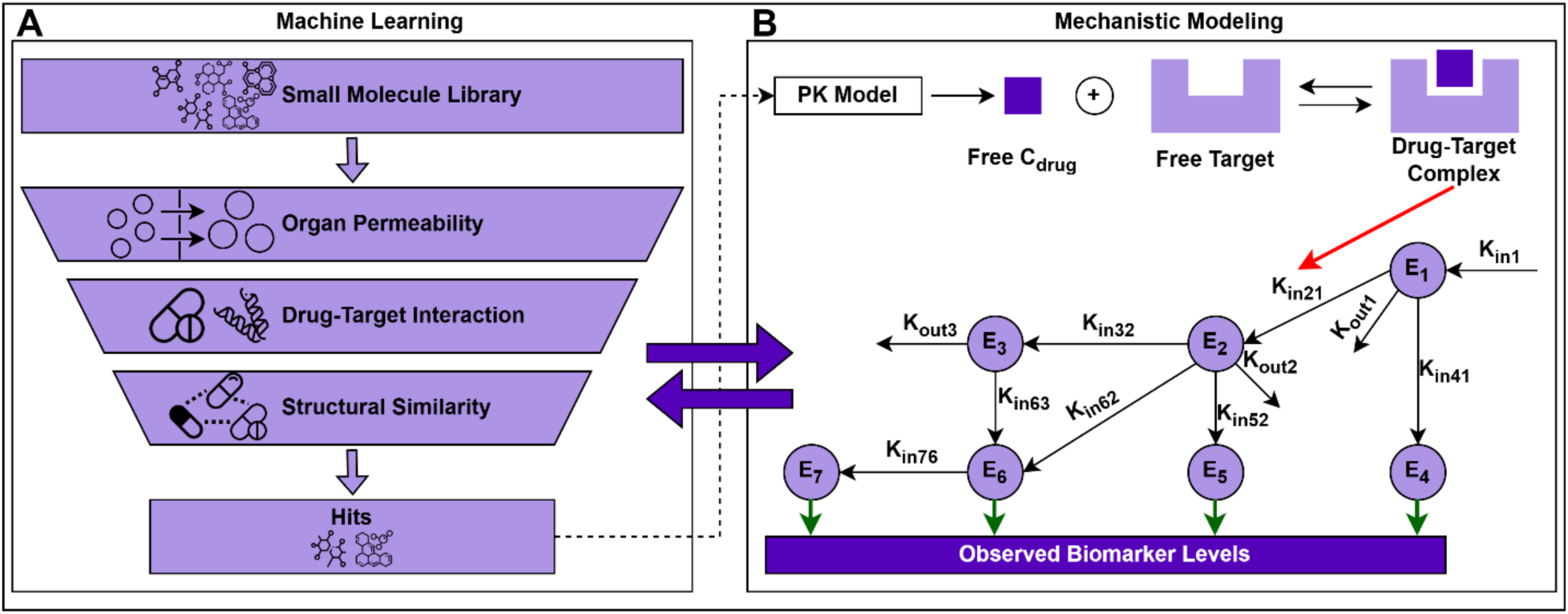
Schematic overview of the proposed framework, Screening Candidates via Organ Uptake and Target-binding (SCOUT). SCOUT integrates **(A)** Machine Learning and **(B)** Mechanistic Modeling. Machine Learning arm screens for candidate hits from a vast small molecule library via three different modules: Organ permeability, Drug-target interaction, and Structural similarity. Mechanistic Modeling arm analyses the potential impact of the candidate hits from machine learning on the downstream modulation of relevant disease biomarkers. The hits from machine learning are used for further analysis via mechanistic model, and the design of mechanistic model governs several choices for parameters in machine learning development, therefore it is a two-way street as shown by double-headed arrows.

The ML arm consists of three sequential modules, (i) organ permeability, (ii) drug-target interaction (DTI) and (iii) structural similarity. Organ permeability predicts the drug’s ability to reach the target organ or site of action (e.g. crossing the BBB). DTI identifies drug candidates capable of binding effectively to the biological target of interest. Structural similarity benchmarks identified candidates against known treatments to identify both chemical novelties and established scaffolds. The schematic of the machine learning arm is shown in Figure 1A.

In this framework, mechanistic modeling is defined as the integration of Pharmacokinetics (PK) and Quantitative Systems Pharmacology (QSP). The PK module estimates the free drug concentration available at the site of action, while the QSP module simulates the dynamic interactions within a disease pathway. By combining these, SCOUT can predict how a drug-target complex modulates downstream biomarker levels (e.g., E4–E7 in Figure 1B), moving beyond simple binding predictions to evaluate functional therapeutic impact.

Critically, the integration of these methodologies is bidirectional. While the ML arm provides promising hits for mechanistic analysis, the mechanistic QSP model informs the initial ML design. Specifically, the disease pathway analysis in the QSP module helps in (i) Target Selection i.e. identifying which proteins in the network are linked to the disease of interest, and (ii) Affinity Metrics i.e. determining whether Ki, IC50 or Kd is the most relevant parameter for the quantifying binding affinity. This “two-way street” design ensures that the ML screening is biologically grounded from the outset, focusing the search space on candidates that are not just chemically viable, but mechanistically relevant to the disease of interest.

Further, confidence in linking machine learning and mechanistic modeling depends on the ability of these models to correctly capture disease biology. The focus of this study is neither to develop a new mechanistic model in-house nor to validate existing ones. The main aim is to develop a framework which integrates machine learning and mechanistic modeling to provide insights for drug repurposing. For PoC, we have focused on small molecule drug repurposing for Alzheimer’s disease in early research or development.

### SCOUT predicts Blood Brain Barrier permeability with significant accuracy

Blood Brain Barrier (BBB) is a specialized semi-permeable membrane responsible for brain’s internal environment and provides protection from harmful foreign chemicals [40]–[43]. For the Alzheimer’s disease PoC, the organ permeability module was designed to predict the BBB permeability. We prioritized the unbound brain-to-plasma partition coefficient (Kp,uu) [23] over traditional blood-brain partition coefficient (logBB), as Kp,uu specifically accounts for the pharmacologically active drug fraction available to bind with neurotherapeutic targets. On the other hand, though logBB has been extensively used, it accounts for both bound and unbound drug concentrations which can be misleading [44].

The organ permeability module demonstrated robust predictive power across both continuous and categorical evaluations. The schematic of the machine learning model used to predict Kp,uu is shown in Figure 2A. In regression analysis, the model achieved a significant Spearman rank correlation of 0.80 (p = 1.61E-20), and coefficient of determination (R^2^ = 0.57) using five-fold cross validation approach, indicating a high degree of monotonic relationship between predicted and observed partition coefficients (Figure 2B). Further, we use the threshold of Kp,uu = 0.3 [22], [45], [46], to evaluate the classification performance of the regression model (classification accuracy: 0.86), as shown in Figure 2C. Using a similar threshold for developing a classification model, the module achieved an accuracy of 90%, recall of 0.94 and an AUC of 0.92 (Figures 2D, 2E). This high sensitivity ensures that the framework can effectively prioritize candidates with sufficient CNS exposure while minimizing false positives. The other relevant regression and classification performance metrics are shown in Table 1.

**Figure 2.**
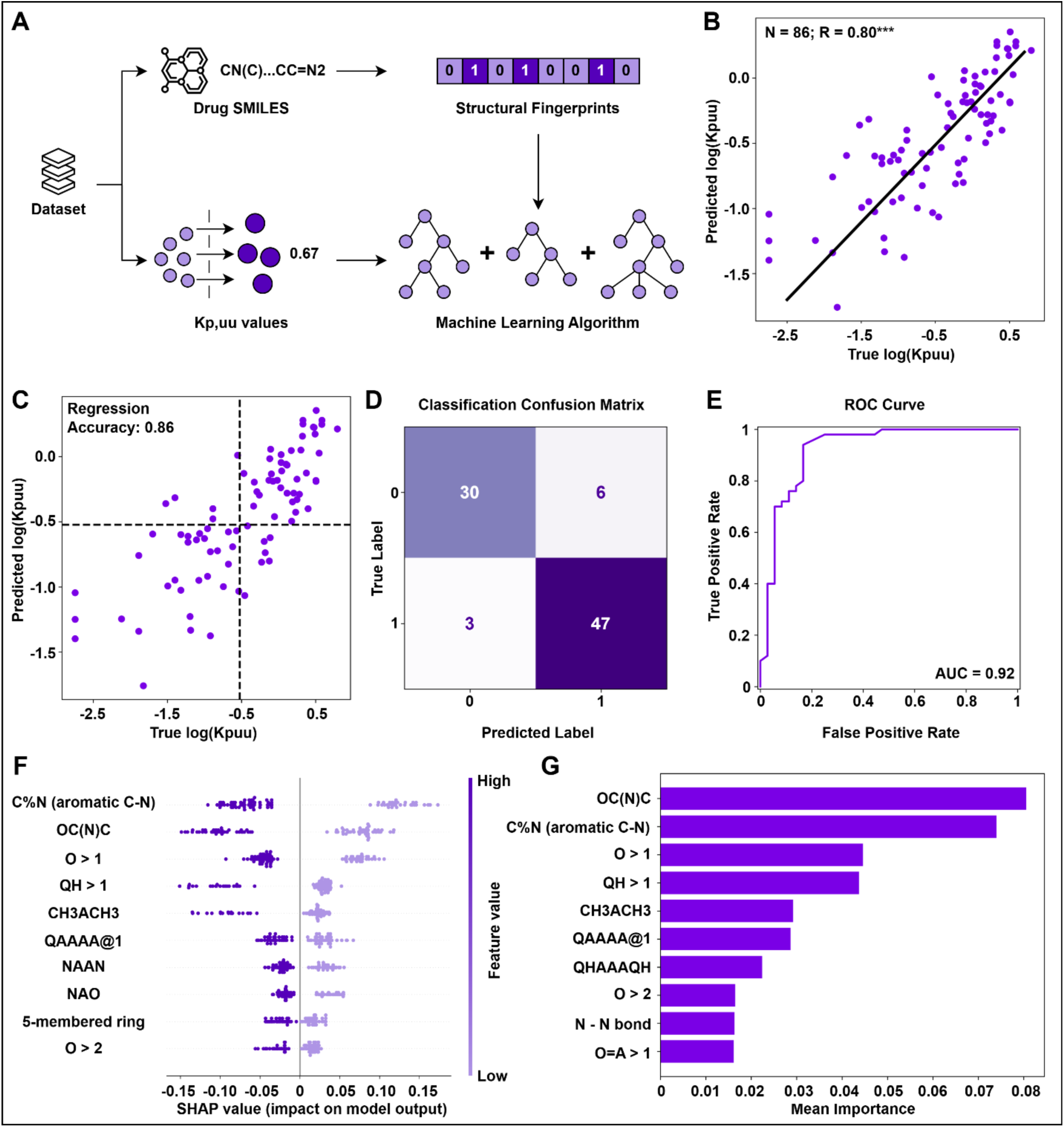
Overview, Performance and Interpretation of Kp,uu model development. **(A)** Schematic overview of Kp,uu model development by utilizing structural fingerprints (MACCS Keys) derived from SMILES notation of drugs. **(B)** Scatterplot show Kp,uu regression prediction model noting number of data points (N), and Spearman rank correlation (R). **(C)** Scatterplot show classification accuracy of Kp,uu regression prediction model by using threshold (Kp,uu = 0.3). **(D)** Confusion matrix show the performance of Kp,uu classification model trained using same threshold as in (C). **(E)** ROC curve shows classification model performance across all thresholds and displaying tradeoff between true positive rate and false positive rate. **(F)** Shapley summary plot displays the top ten important features and impact of their presence or absence on individual data points. **(G)** Bar plot displays the top ten important features using Permutation importance, based on the impact on overall model performance when a particular feature is randomly shuffled.

**Table 1.** Performance metrics for Regression and Classification models for Kp,uu prediction.

|  |  |
| --- | --- |
| Regression Performance Metrics |  |
| Spearman correlation (R) | 0.80 |
| P-value | 1.60E-20 |
| Pearson correlation | 0.79 |
| Mean Absolute Error (MAE) | 0.41 |
| Mean Squared Error (MSE) | 0.29 |
| Coefficient of Determination (R2) | 0.57 |
| Absolute Fold Error | 2.57 |
| Geometric Fold Error | 1.03 |
| Median Fold Error | 2.42 |
| Regression-based Classification Metrics |  |
| Accuracy | 0.86 |
| F1-score | 0.88 |
| Sensitivity | 0.86 |
| Matthews Correlation Coefficient (MCC) | 0.72 |
| Area Under Curve - ROC | 0.86 |
| Classification Performance Metrics |  |
| Accuracy | 0.90 |
| F1-score | 0.91 |
| Sensitivity | 0.94 |
| Matthews Correlation Coefficient (MCC) | 0.78 |
| Area Under Curve - ROC | 0.92 |

The module performance compares favorably against established benchmarks in literature. While previous models using molecular descriptors or solvation energy achieved accuracy ranging from 79% [26] to 85% [24], SCOUT’s classification accuracy of 90% represents a significant improvement (Table 1). Furthermore, SCOUT achieves these results using a streamlined structural input, avoiding the need for high-cost graph convolutional network (GCN) approach that achieved R^2^ of 0.36 [25]. We attribute this improved performance to a combination of rigorous data preprocessing specifically the log-transformation of Kp,uu values to normalize variance, extensive hyperparameter optimization, and the strategic use of rule-based chemical embeddings that capture essential structural motifs more effectively than raw descriptors. Central to the broad utility of these predictions is the assumption that Kp,uu values measured in rodents are representative of those in primates, including monkeys and humans. This cross-species translation is supported by the high evolutionary conservation of BBB physiology and transporter-mediated efflux mechanisms across various species [47]–[49]. These results establish the organ permeability module as a high-precision first-tier filter within the SCOUT pipeline, successfully de-risking the search space for BBB permeable compounds.

### Model Interpretation analysis validates the mechanistic consistency of the Kp,uu model

To ensure that the high predictive accuracy of the organ permeability module is grounded in established chemical principles, we performed an interpretability analysis using SHAP (SHapley Additive exPlanations) values [50] and permutation importance [51] (Figures 2F and 2G). This dual-method approach serves as a critical sanity check, validating that the model has focused on the physical laws governing blood-brain barrier (BBB) penetration rather than relying on spurious correlations, for making predictions.

The convergence of both SHAP and permutation importance identified five primary structural determinants of Kp,uu prediction: (i) Aromatic nitrogen (C%N), whose presence increases molecular polarity and polar surface area (PSA), (ii) Amide-like configurations (OC(N)C), which increases local dipole moments due to proximity of oxygen and nitrogen on a single carbon, (iii) Oxygen density (O>1), that increases hydrogen-bond acceptor (HBA) potential, (iv) Heteroatom-Hydrogen bonding (QH>1), indicating the presence of multiple polar groups (O, N, S, P) capable of hydrogen bonding, and (v) Methyl branching (CH3ACH3) reflecting molecular bulk and overall size.

The model’s internal logic remains highly consistent with the known pharmacokinetics of CNS drugs i.e. higher values for these polar features (increased PSA and H-bond potential) contributed significantly to a decrease in predicted Kp,uu (Figure 2F) [52], [53]. This alignment with established medicinal chemistry rules, specifically that increased polarity and molecular size impede passive diffusion across the BBB, validates the reliability of the SCOUT framework. Rather than acting as a black box model, the permeability module functions as a transparent predictive tool that accurately captures the structural features governing drug distribution in the brain.

### SCOUT utilizes structural features of drugs and targets to predict drug-target interactions

Small molecule drugs exert their therapeutic influence via perturbation of protein networks [54], therefore modeling drug-protein interactions is crucial for drug repurposing. Following the identification of brain-permeable candidates, the SCOUT framework employs a Drug-Target Interaction (DTI) module, as conducted in a recent study [55], to predict binding efficacy (Figure 3A). For this Alzheimer’s disease PoC, we focused on BACE1 (β-secretase 1), a well-characterized enzyme responsible for the initial step in Aβ production [27]–[30].

**Figure 3.**
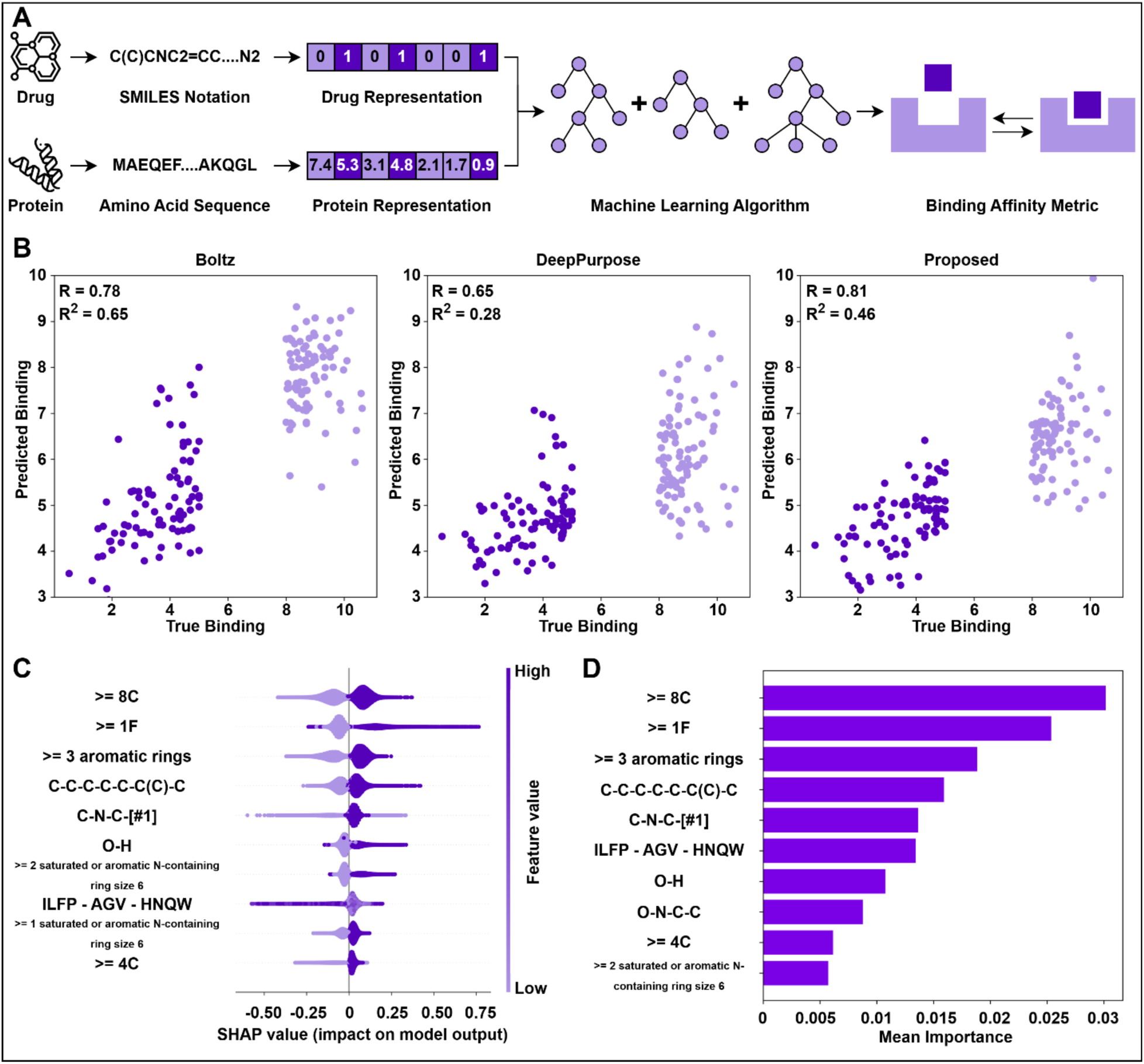
Overview, Performance and Interpretation of DTI model development. **(A)** Schematic overview of DTI model development by utilizing structural fingerprints of both drugs and proteins derived from SMILES notation and Amino acid sequence respectively. **(B)** Scatterplots showing Spearman rank correlation and Coefficient of determination for true and predictive binding for benchmarking proposed approach (SCOUT) with literature approaches (Boltz-2, and DeepPurpose). **(C)** Shapley summary plot displays the top ten important features and impact of their presence or absence on individual data points. **(D)** Bar plot displays the top ten important features using Permutation importance, based on the impact on overall model performance when a particular feature is removed.

A unique strength of the SCOUT framework is the bidirectional flow of information between machine learning and mechanistic modeling arms. While our ML arm can predict multiple affinity metrics, including Kd, Ki, and EC50, our downstream QSP model specifically requires IC50 values to simulate the kinetic formation of the drug-target complex. Consequently, we prioritized the development and validation of the pIC50 (negative log of IC50 values) predictive model to ensure seamless integration for the final mechanistic simulation.

The DTI module demonstrated high predictive accuracy for predicting pIC50. The regression model achieved a Spearman rank correlation of 0.86 (p ∼ 0), indicating a high level of agreement between predicted and experimental binding values (Table 2). To evaluate its utility as a binary screening tool, we applied a threshold of pIC50 > 6, i.e. IC50 < 1000 nM. In this classification task, the module achieved an accuracy of 87% and an exceptionally high AUC of 0.95 (Table 2). These metrics confirm that the DTI module can accurately distinguish between potent binders and weak binders, providing a robust second-stage filter that ensures only highly active compounds are selected for mechanistic validation.

**Table 2.** Performance metrics for Regression and Classification models for DTI prediction (pIC50).

|  |  |
| --- | --- |
| **Regression Performance Metrics** |  |
| Spearman correlation (R) | 0.86 |
| P-value | ~0 |
| Pearson correlation | 0.86 |
| Mean Absolute Error (MAE) | 0.57 |
| Mean Squared Error (MSE) | 0.59 |
| Coefficient of Determination (R2) | 0.74 |
| **Regression-based Classification Metrics** |  |
| Accuracy | 0.86 |
| F1-score | 0.87 |
| Sensitivity | 0.87 |
| Matthews Correlation Coefficient (MCC) | 0.72 |
| Area Under Curve - ROC | 0.86 |
| **Classification Performance Metrics** |  |
| Accuracy | 0.87 |
| F1-score | 0.88 |
| Sensitivity | 0.89 |
| Matthews Correlation Coefficient (MCC) | 0.75 |
| Area Under Curve - ROC | 0.95 |

### SCOUT is benchmarked for predicting drug-target interaction scores against state-of-the-art predictive architectures

To evaluate the predictive performance of the DTI module within the context of high-throughput screening, we benchmarked SCOUT against two established methodologies: DeepPurpose, a deep-learning framework utilizing sequence-based featurization [34], and Boltz-2, a state-of-the-art Large Language Model (LLM) that utilizes 3D structural data for binding affinity prediction [35]. Given the extreme computational requirements of 3D-aware modeling in Boltz-2, which can be prohibitive for screening massive chemical libraries, we conducted this comparative analysis on a representative subset of ‘good’ and ‘bad’ binders (Figure 3B).

The results demonstrate that SCOUT achieves a significant degree of predictive alignment with true binding scores. Within this focused evaluation, SCOUT displayed superior performance metrics compared to DeepPurpose and reached performance levels comparable to Boltz-2. While Boltz-2 offers significant technological advancements in 3D-structural awareness, it encountered structural processing errors for a small subset of the tested pairs; in contrast, SCOUT provided robust, uninterrupted predictions across the entire dataset.

These findings suggest that for the specific task of high-throughput screening in drug repurposing, where speed and scalability are as critical as precision, SCOUT serves as a highly effective fit-for-purpose framework. By utilizing optimized chemical embeddings rather than resource-intensive 3D-structural data, SCOUT provides the high-throughput reliability necessary to filter 20,000+ compounds without the prohibitive computational overhead associated with 3D structural LLMs.

### Interpretability Analysis Validates DTI Model Alignment with Medicinal Chemistry Principles

To ensure the DTI module provides reliable predictions, we utilized SHAP values and permutation importance to examine the internal logic of the model (Figures 3C and 3D). This analysis serves to validate that the framework identifies candidates based on established biochemical drivers of affinity rather than algorithmic artifacts.

The model’s interpretation revealed that its top predictive features align closely with the hydrophobic drive, commonly observed in potent small-molecule binders. Features associated with carbon skeletons and aromatic rings, such as “>= 8 carbons”, “>= 3 aromatic rings” and long-chain alkanes, were the strongest positive predictors of binding affinity (Figure 3C). This demonstrates that the model correctly prioritized the hydrophobic effect, where lipophilic drug motifs bind effectively into the hydrophobic pockets of protein targets [56]–[59].

Furthermore, the model identified fluorination (“>= 1F”) as a critical contributor to increased binding affinity. In medicinal chemistry, fluorine is strategically used to strengthen multipolar interactions and lock molecules into their bioactive conformations [60]–[62]. SCOUT’s identification of this fluorine effect as a top contributor significantly bolsters confidence in its predictive logic. Additionally, the model correctly weighed the presence of hydrogen bond donors and acceptors, e.g., hydroxyl groups (“O-H”) and secondary amines (“C-N-C-[#1]”), identifying their role in forming directional bonds with specific protein residues [63].

The model further identified the presence of multiple Nitrogen-containing rings as a key driver of high binding affinity. This observation aligns with the established role of basic nitrogenous motifs in forming salt bridges and stabilizing interactions with target residues. Notably, the importance of this feature reflects the model’s ability to recognize structural requirements common in successful small-molecule drugs, such as Kinase inhibitors, where N-containing scaffolds are ubiquitous for providing structural stability and directional binding [64]–[69]. By prioritizing these motifs, SCOUT demonstrates an understanding of the electrostatic complementarity required for potent target inhibition.

The interpretability analysis also extended to the protein sequence, identifying a specific amino acid triad: [ILFP]–[AGV]–[HNQW], as a key determinant of the binding affinity. The model identified this specific sequence as a strong negative predictor of affinity. Biologically, this triad represents a transition from large hydrophobic residues to a polar patch. The model’s determination that this environment is poorly compatible with drug binders suggests it has successfully captured the risk of steric clashes or unfavorable electrostatic interactions within certain protein regions [70], [71].

By identifying these key structure-activity relationships (SAR), the SCOUT framework demonstrates that its predictions align with fundamental biochemical principles. While the underlying machine learning architecture retains its inherently complex ‘black-box’ nature, the recovery of these patterns provides a layer of mechanistic plausibility by highlighting the molecular motifs that drive high-affinity binding. Although drug-target interaction remains an open and mathematically challenging problem, the model’s ability to capture these biological sensitivities suggests that it is leveraging meaningful chemical signals rather than arbitrary noise. These interpretative insights serve to rationalize the model’s outputs, offering researchers a more grounded basis for prioritizing candidates during the early stages of drug discovery.

### Analysis of Kp,uu-DTI landscape for the Broad Repurposing Hub and Drugbank libraries

As SCOUT has shown significant accuracy and performance in predicting Kp,uu and DTI scores, we integrated the outputs of these modules into a Kp,uu-DTI Landscape. This dual-parameter scatterplot represents Kpuu scores (log(Kp,uu)) along y-axis and DTI scores (pIC50) along x-axis, with each point corresponding to a specific drug in the small molecule library. The candidate hits will be positioned in the top-right quadrant, where drugs possess both high predicted brain penetration and high binding affinity for BACE1. By evaluating these metrics simultaneously, SCOUT acts as a co-optimization filter, identifying the compounds with target potency as well as organ-level access (Figure 4A).

**Figure 4.**
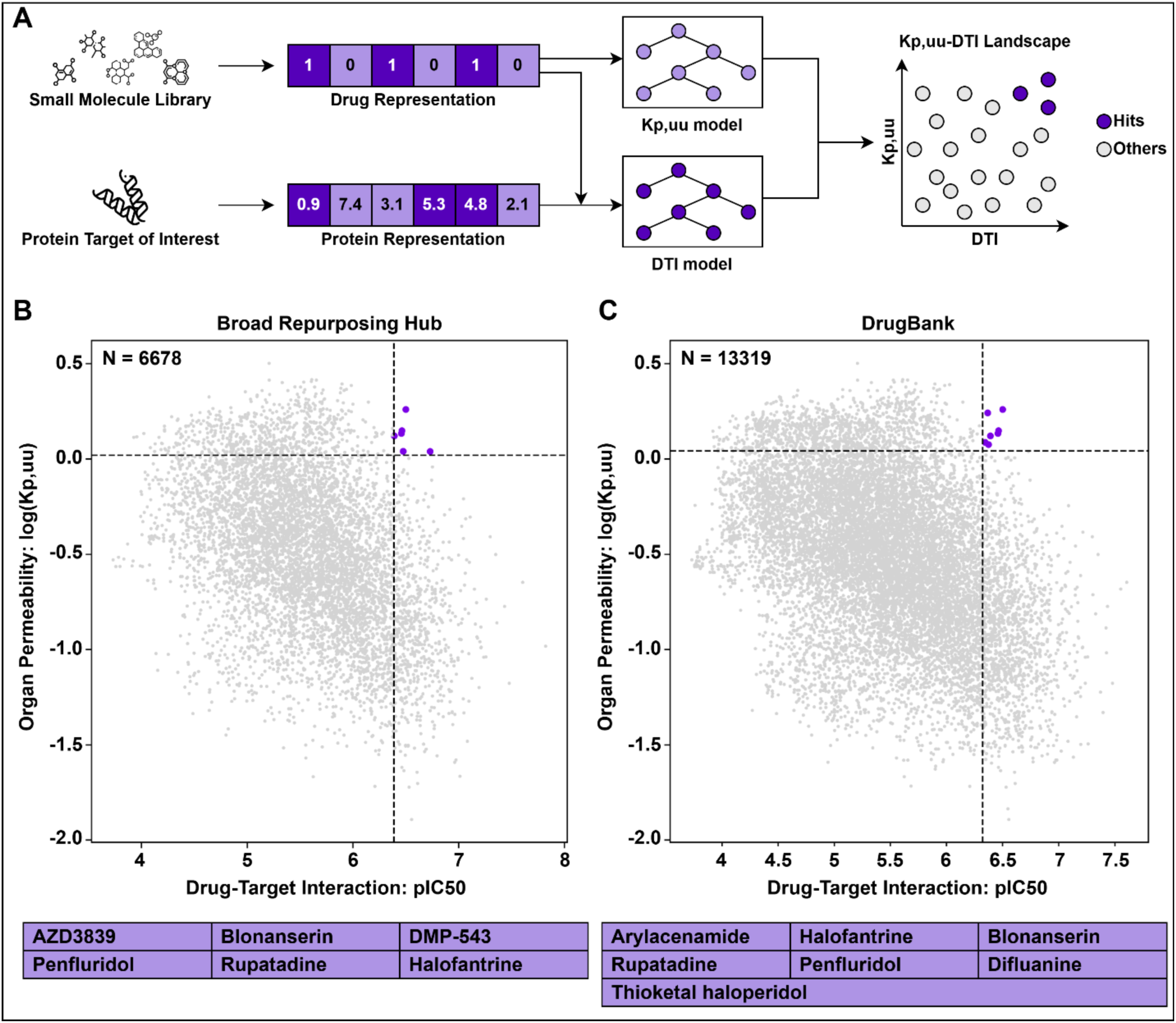
Kp,uu-DTI Landscapes for Screening Libraries. **(A)** Schematic overview of generating a Kp,uu-DTI Landscape using both Kp,uu and DTI predictive models. Compounds in the library with high Kp,uu and DTI metrics are highlighted in purple. Kp,uu-DTI Landscape for **(B)** Broad Repurposing Hub library (6678 compounds) and **(C)** DrugBank small molecule library (13319 compounds).

We applied the landscape analysis to two comprehensive small-molecule libraries: the Broad Repurposing Hub [72] and DrugBank [73]. To identify the most promising candidates for downstream validation, we implemented a stringent prioritization strategy targeting compounds within the top 10% for both predicted permeability (Kp,uu) and binding affinity (DTI). While a random distribution would theoretically yield a 1% overlap, our framework identified significantly more refined hits: only 6 compounds from the Broad library (0.09%) and 7 compounds from DrugBank (0.05%) met these dual criteria (Figures 4B and 4C). Rather than serving as a direct metric of model accuracy which was validated in the preceding individual modules, this outcome highlights the framework’s utility as a stringent decision-support tool. The low hit rate observed reflects the inherent scarcity of ‘dual-optimized’ hits that satisfy the requirements for both CNS penetration and target potency. By narrowing the experimental search space by nearly 99.9%, the SCOUT framework does not solve the discovery problem but rather provides a highly prioritized and resource-efficient roadmap. This allows researchers to focus secondary, high-cost validation efforts on a statistically elite subset of candidates with the highest probability of satisfying the multi-parameter requirements.

The candidate hits identified by SCOUT represent a diverse pharmacological landscape, spanning several development stages and disease indications (Table 3). The observed hits include both established AD-related agents and compounds with mechanisms of action that have been independently investigated in the context of neurodegeneration.

**Table 3.** The list of compounds hits identified by SCOUT framework from Broad repurposing hub and Drugbank small molecule screening libraries, along with information on phase in pipeline, mechanism, disease area (indication), brief description, predicted Kp,uu and IC50 scores, and computed tanimoto similarity scores using MACCS keys and Pubchem substructure fingerprints. Drugbank ID in parenthesis. *indicates compound hit identified in both libraries.

|  |  |  |  |  |  |  |  |  |
| --- | --- | --- | --- | --- | --- | --- | --- | --- |
| Drug | Phase | Mechanism | Disease Area (Indication) | Description | Kp,uu | IC50 (uM) | MACCS | Pubchem |
| AZD3839 | Phase 1 | BACE1 inhibitor | - | Inhibits the BACE1 enzyme, responsible for cleaving Amyloid Precursor Protein (APP), preventing the production of Amyloid-beta proteins before they aggregate into plaques. | 1.09 | 0.19 | 0.343 | 0.545 |
| Blonanserin* (DB09223) | Launched | Dopamine receptor antagonist<br>Serotonin receptor antagonist | Neurology/<br>Psychiatry (Schizophrenia) | Antagonist for Dopamine D2 and D3, and Serotonin 5-hydroxytryptamine receptor 2A. Shows high affinity for D2 receptor controlling positive symptoms like hallucinations and delusions. Blocks serotonin receptor to help with negative symptoms like social withdrawal or lack of motivation. | 1.36 | 0.35 | 0.337 | 0.555 |
| DMP-543 | Phase 2 | Acetylcholine release enhancer | - | Blocks Kv7.2 (KCNQ2) potassium channel and enhances the release of Acetylcholine, a primary neurotransmitter. Increases the release of Dopamine and Glutamate. Possesses more fluorine atoms than its parent compound, Linopirdine providing more metabolic stability. | 1.1 | 0.33 | 0.284 | 0.587 |
| Halofantrine* (DB01218) | Launched | Voltage-gated inwardly rectifying potassium channel KCNH2 inhibitor | Infectious disease (Malaria) | Kills the malarial parasite during its stage in red blood cells. Does not prevent malaria and is ineffective against parasite stages in the liver. | 1.32 | 0.41 | 0.278 | 0.364 |
| Penfluridol* (DB13791) | Launched | T-type calcium channel blocker | Neurology/<br>Psychiatry (Schizophrenia) | Antagonist for Dopamine D2 receptor, with additional effects on Serotonin, and Histamine. Shows exceptionally long half-life leading to a weekly dose. | 1.41 | 0.34 | 0.293 | 0.383 |
| Rupatadine*<br>(DB11614) | Launched | Histamine<br>receptor<br>antagonist<br>Platelet activating<br>factor receptor<br>antagonist | Allergy<br>(Allergic rhinitis,<br>Urticaria) | Non-sedating antihistamine. Both the<br>receptors are involved in inflammatory<br>responses. | 1.82 | 0.32 | 0.27 | 0.509 |
| Difluanine<br>(DB19523) | Experimental | Dopamine<br>receptor<br>antagonist | - | Antagonist for L-type calcium channels.<br>Acts as CNS stimulant. | 1.22 | 0.45 | 0.296 | 0.56 |
| Arylacenamide<br>(DB05792) | Experimental | Sigma receptor<br>ligand<br>Steroid<br>isomerase<br>inhibitor |  | Binds four proteins in human cells, SRBP-1, [sigma]-2, HSI, and its relative SRBP-2.<br>Dual agent with both immunomodulatory<br>and antiproliferative activities | 1.74 | 0.43 | 0.23 | 0.382 |
| Thioketal<br>haloperidol<br>(DB08622) | Experimental | Dopamine<br>receptor<br>antagonist |  | Modulate dopamine D2 receptors or sigma<br>receptors. Thioketal group is inactive in<br>normal conditions. Under high oxidative<br>stress, molecules break into normal<br>haloperidol, binding with dopamine<br>receptors. | 1.19 | 0.42 | 0.31 | 0.404 |

The framework successfully identified AZD3839, a known BACE1 inhibitor [74], providing a direct internal validation of the DTI module’s accuracy. Additionally, the pipeline prioritized Donepezil, an FDA-approved treatment for AD-related dementia that functions as an acetylcholinesterase inhibitor [75], [76]. Beyond these established agents, SCOUT highlighted DMP-543, an acetylcholine release enhancer. The prioritization of DMP-543 is consistent with research exploring the augmentation of synaptic acetylcholine supply to address the cholinergic deficits characteristic of AD pathology [77]–[79]

Several identified hits correlate with pharmacological strategies aimed at addressing the neuronal hyperexcitability [80]–[82] and calcium dysregulation [83], [84] reported in early-stage AD. Penfluridol, a calcium channel blocker, appeared within the high-priority hits. Such compounds have been studied in independent research for their potential to stabilize neuronal firing [85]. Similarly, Halofantrine, which has been reported to modulate KCNH2 (hERG) channels, aligns with theoretical frameworks investigating the dampening of excitotoxicity-induced cognitive decline [86].

SCOUT identified several dopamine, serotonin and histamine receptor antagonists, Blonanserin, Rupatadine, Difluanine, and Thioketal haloperidol, typically used in psychiatric indications [87], [88]. While these could be proposed for managing the neuropsychiatric symptoms of dementia (agitation and aggression), they may also possess disease-modifying potential. For instance, dopamine D2 antagonism is hypothesized to reduce Tau aggregation [89], while specific serotonin receptor (5-HT6) modulation is being investigated for its potential to improve cognitive function [90], [91].

A subset of the identified candidates targets pathways linked to cellular resilience against Aβ toxicity. Arylacenamide, prioritized as a putative Sigma receptor modulator, has been discussed in literature regarding protein folding, mitochondrial protection, and the mitigation of endoplasmic reticulum (ER) stress, which is associated with misfolded Aβ and phosphorylated Tau [92]–[96]. Rupatadine, a Platelet Activating Factor (PAF) receptor antagonist, has also been studied in the context of neuroinflammation and the maintenance of synaptic integrity [97]–[99]. Furthermore, the prioritization of antimalarial agents such as Halofantrine coincides with research into autophagy enhancement as a potential mechanism for the clearance of toxic protein aggregates [100], [101].

It is important to note that these biological associations are provided as a post-hoc analysis of the model’s output and do not imply a definitive therapeutic claim. As a computational tool, SCOUT provides a structured prioritization based on predicted binding and permeability; however, the functional relevance of these hits requires prospective experimental validation. These results serve to demonstrate that the framework’s high-priority outputs are grounded in recognized pharmacological principles, distinguishing meaningful chemical signals from computational noise.

### Structural comparison of candidate hits with known inhibitors provides a confidence metric

The SCOUT framework utilizes the Similar Property Principle (SPP), the concept that structurally related molecules often share similar physicochemical and biological activities [102], to benchmark candidate hits against established BACE1 inhibitors. By comparing our identified hits to known clinical-stage inhibitors as per Drugbank i.e. Verubecestat (DB12285), CTS-21166 (DB06073), and Elenbecestat (DB15391), we can distinguish between similar candidates that resemble proven scaffolds and novel candidates that expand the targetable chemical space.

We quantified structural similarity using Tanimoto coefficients [103] calculated across two distinct fingerprinting methods. Our results indicate that PubChem substructure fingerprints provide significantly higher granularity for candidate differentiation compared to MACCS keys (Table 3). While MACCS keys yielded relatively uniform scores across all hits, likely due to the limited 166-bit resolution of the fingerprinting method, the more detailed PubChem fingerprints (881-bit resolution) revealed a clear stratification of candidates.

Candidates with primary mechanisms involving neural activity either by inhibiting BACE1 (AZD3839) or involved in modulating neurotransmitters (Blonanserin, DMP-543, Rupatadine, Difluanine, and Thioketal haloperidol) exhibited higher similarity to known BACE1 inhibitors. This structural overlap provides a confidence metric, suggesting these drugs possess the requisite chemical motifs for effective BACE1 binding. Conversely, several candidates showed low structural resemblance to established BACE1 inhibitors. These divergent scaffolds are of particular interest as they may offer opportunities to avoid the cross-reactivity or class-specific side effects associated with current BACE1 therapeutics. By integrating this structural similarity module, the SCOUT framework not only just output a list of drugs but also provides a rational basis for selecting candidates that balance mechanistic validation (high similarity) with therapeutic novelty (low similarity).

### Mechanistic modeling establishes a framework for analyzing the impact of compound hits on modulation of downstream biomarkers

While standard ML-based drug repurposing approaches identify promising candidates based on binding probability, SCOUT extends this analysis by simulating the modulation of disease biomarkers. By integrating the ML-predicted Kp,uu and IC50 values into a mechanistic QSP framework, we bridge the gap between biological affinity and functional therapeutic efficacy (Figure 5A).

**Figure 5.**
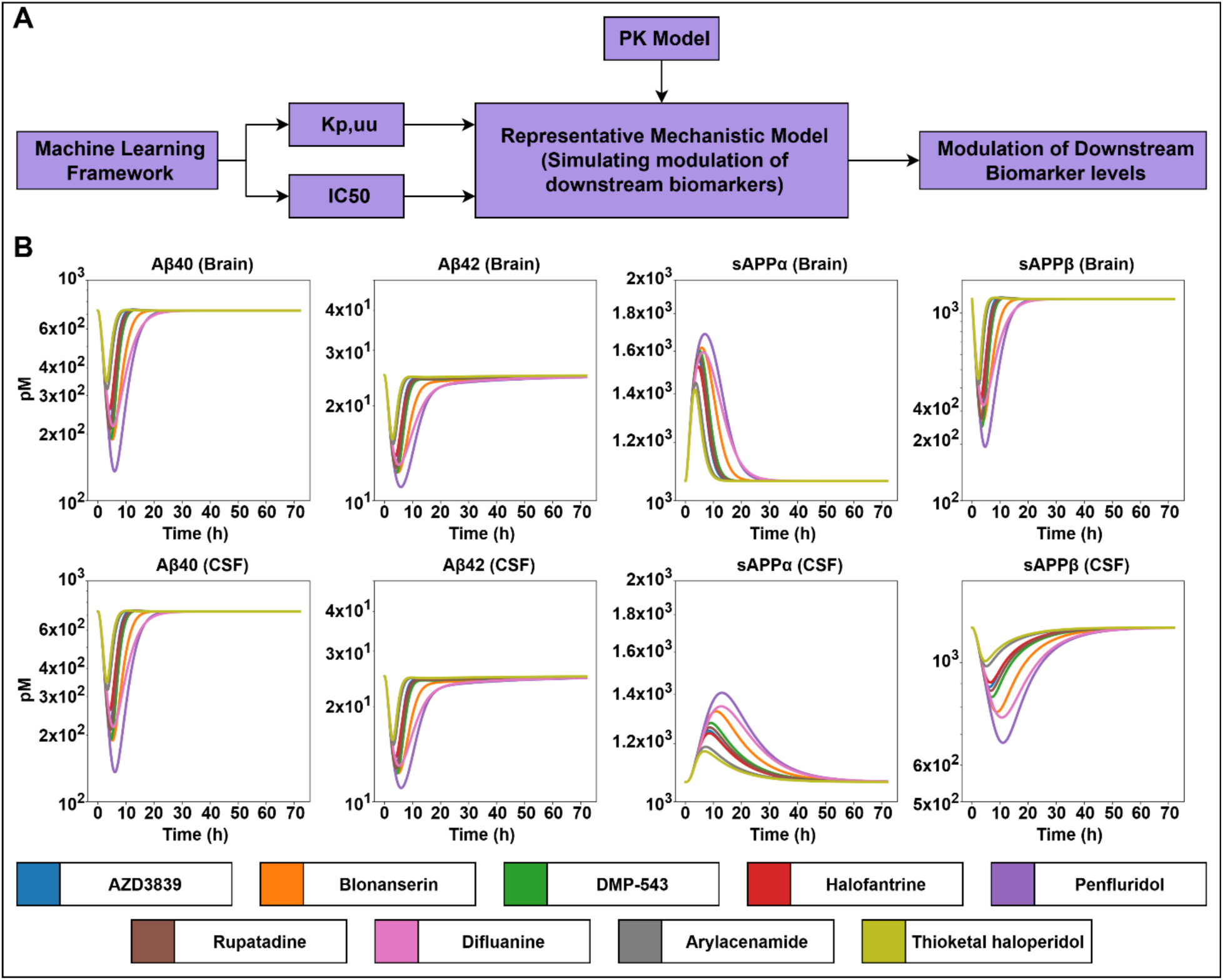
Mechanistic model simulates modulation in downstream biomarkers for compound hits. **(A)** Schematic overview of combining predicted scores of Kp,uu and IC50 from machine learning models along with PK model to simulate modulation of downstream biomarker levels using representative mechanistic model. **(B)** The effect of each compound hit on the modulation of all four different biomarkers, in both brain and CSF region. Y-axis in all the plots have units of pM (picomolar).

For this PoC, we utilized a translationally validated systems pharmacology model developed by Leiden and Merck [39]. While originally parameterized using Rhesus monkey data, this model serves as a robust kinetic surrogate for human Aβ dynamics, as the amyloidogenic pathway and BACE1 target homology are highly conserved among primates [104]–[106]. By feeding our ML-derived scores (Kp,uu and IC50) of the candidate hits along with PK parameters (clearance and volume of distribution) obtained from the model previously published [107] into this QSP model (Supplementary Table 10), we transformed static hits into dynamic simulations of disease modulation.

It is important to acknowledge that the integration of multiple machine learning modules, predicting permeability, binding affinity, and pharmacokinetics introduces the risk of compounded predictive error. Uncertainties inherent in each individual ML model can propagate and potentially amplify within the mechanistic simulation. However, we emphasize that in real-world drug repurposing applications, the pharmacokinetic profiles of these candidates would typically be known from established preclinical or clinical datasets. In such use-cases, these experimental PK values would replace the ML-based estimates used in this PoC, significantly refining the precision of the downstream biomarker simulations and mitigating the risk of error propagation.

The QSP simulations revealed a consistent mechanistic shift across all candidate hits. As shown in Figure 5B, the inhibition of BACE1 successfully diverted Amyloid Precursor Protein (APP) processing from the amyloidogenic pathway to the non-amyloidogenic pathway. This was evidenced by a significant reduction in neurotoxic biomarkers Aβ40, Aβ42, and sAPPβ) and a concomitant increase in the neuroprotective sAPPα fragment.

A major benefit of this modular approach is the ability to rank-order candidates based on predicted clinical endpoints. For instance, while several hits showed similar DTI binding scores, their differing Kp,uu (permeability) values led to varied levels of Aβ reduction in the brain compartment. This allows for a more rational selection of leads that optimize both potency and site-of-action concentration.

To validate the necessity of our multi-stage funnel, we simulated non-hit compounds, those with poor predicted permeability or weak binding affinity. These negative controls failed to produce significant biomarker modulation (data shown in Supplementary Figures 3 and 4), confirming that the ML arm effectively pre-screens only those candidates capable of driving a meaningful biological response. This integration ensures that SCOUT provides a resource-efficient pathway, focusing experimental efforts only on compounds with the highest probability of functional success.

## Discussion

SCOUT represents a shift in computational drug repurposing, moving away from fragmented black box predictions toward an explainable, modular workflow architecture. By connecting structural SMILES and amino acid sequences directly to functional Aβ biomarker modulation, the framework provides a multi-stage de-risking strategy for neurological drug development.

The primary success of the SCOUT framework lies in its ability to resolve the needle-in-a-haystack problem that plagues traditional screening. By co-optimizing organ permeability (Kp,uu) and target affinity (IC50), we achieved a 99.9% reduction in the candidate search space. For a pharmaceutical program, this translates to a massive reduction in preclinical timelines and costs. Rather than conducting broad, low-probability animal experiments, researchers can utilize SCOUT to prioritize a top-tier list of candidates. This approach directly supports the 3Rs (Replacement, Reduction, and Refinement) of animal testing and aligns with the strategic direction of the FDA Modernization Act 2.0. By providing a robust computational surrogate for initial efficacy screening, SCOUT fits within the emerging category of New Approach Methodologies (NAMs) that the FDA and other global regulators are increasingly encouraging to phase out unnecessary animal models. Consequently, the framework ensures that in vivo resources are reserved only for candidates with the highest probability of clinical success.

A key consideration in the development of the SCOUT framework was the selection of the DTI predictive architecture. While state-of-the-art 3D-aware models, such as Boltz-2, offer high-resolution insights into binding orientations, their deployment in high-throughput repurposing campaigns is often limited by significant computational overhead. For this study, we opted for a streamlined ML approach to prioritize operational scalability. By utilizing sequence-based and rule-based chemical embeddings, SCOUT enables the rapid screening of tens of thousands of compounds, a task that would be computationally prohibitive for 3D structure-based LLMs. Furthermore, our approach ensures model robustness, providing uninterrupted predictions across diverse chemical libraries where 3D conformer generation might otherwise encounter processing errors. Thus, SCOUT serves as a ‘fit-for-purpose’ engine, providing the high-precision filtering necessary to condense vast chemical spaces into manageable, high-priority leads for downstream mechanistic validation.

Another critical consideration in any multi-stage framework is how uncertainty propagates from the machine learning arm to the mechanistic simulation. In this work, we utilize two ML models to feed quantitative parameters into a QSP model. While the ML models showed high Spearman correlations (0.80–0.86), they are not without inherent error. The value of the SCOUT modular approach is that it allows for sensitivity analysis at the simulation stage. We may observe that biomarker levels are not equally sensitive to all inputs i.e. for instance, a 10% error in predicted IC50 may have a negligible impact on Aβ reduction if the drug’s brain permeability (Kp,uu) is the limiting factor. By integrating these modules, we can identify which parameters require the highest experimental precision and which can tolerate the noise typical of in vitro datasets. This transparency allows researchers to quantify the confidence interval of a repurposing hypothesis before moving to the clinic.

The identification of AZD3839 highlights a fundamental challenge in early-stage repurposing: the distinction between a mechanistic hit and a safe drug. While SCOUT correctly identified AZD3839 as a potent BACE1 inhibitor with significant CNS penetration, the drug ultimately failed in Phase 1 trials due to QTc prolongation, a safety issue unrelated to its primary target [74]. This underscores that the ideal application of SCOUT is the screening of established and safe clinical assets (such as Rupatadine or Donepezil). For these drugs, the toxicity profile is already well-documented, significantly reducing the risk of the “unforeseen” failures seen with de novo molecules. Future iterations of SCOUT could mitigate this by incorporating safety-specific ML modules, e.g., hERG inhibition or DILI (Drug-Induced Liver Injury) prediction to further filter candidates before the mechanistic modeling stage.

SCOUT was designed as a plug-and-play architecture to avoid the rigidity of previous repurposing tools. Because each module, Permeability, DTI, Similarity, and QSP is independent, the framework is agnostic to both the disease and the modeling method. Researchers can swap literature-based QSP models for proprietary in-house models or replace tree-based ML with Large Language Models (LLMs) as data availability evolves. While this study focused on the Kp,uu-BACE1 axis in Alzheimer’s disease, the logic is readily extensible to neurodegenerative diseases such as Parkinson’s, Multiple Sclerosis, or even oncology. By providing a scalable, interpretable, and mechanistically grounded roadmap, SCOUT offers a path toward more rational, data-driven drug repurposing that bridges the gap between computational probability and clinical reality.

In summary, the SCOUT framework provides a transformative approach to drug repurposing by bridging the gap between structural information and systems pharmacology. By effectively refining a vast search space of over 10,000 compounds into a focused cohort of high-priority candidates, we demonstrate the framework’s utility as a stringent decision-support tool for the efficient allocation of experimental resources. The subsequent quantification of biomarker modulation transforms these candidates from static structural hits into dynamic therapeutic hypotheses. The modularity of this Two-Way Street between machine learning and mechanistic modeling ensures that SCOUT is not merely a static tool, but an evolving architecture capable of accelerating therapeutic discovery across diverse and complex disease landscapes. Ultimately, this framework offers a resource-efficient pathway to de-risk the transition from preclinical hypotheses to clinical reality, potentially reducing the time and cost associated with bringing therapies to the patients who need them most.

## Methods

### Kpuu Model Development

#### Dataset

The dataset to train Kpuu machine learning model is sourced from the research article by Lawrenz *et al* [26]. The article provides published dataset of Kpuu values for 86 compounds in rat and 37 compounds in mouse. The Kpuu is defined using the equation below,

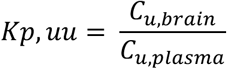

where, *C_u,brain_* and *C_u.plasma_* refer to unbound concentration of drug in brain and plasma respectively. Along with Kpuu values, the dataset provides information on SMILES notation of different compounds, which is an alphanumeric text-based code representing a compound’s chemical structure.

#### Preprocessing

Kp,uu values of different compounds can have different values of magnitude and leads to skewness in the dataset, therefore, we preprocess the Kp,uu values via log transformation (base 10). The transformation helps ensure that there is less skewness in the dataset, and machine learning algorithms can inspect for important patterns rather than failing due to skew data. We have utilized log-transformed values to develop regression model. Further, Kp,uu > 0.3 is considered as an appreciable transport and penetration in rats, so we have used it as a threshold to train classification models.

#### Featurization

As mentioned before, compounds are represented by SMILES notation in the dataset. The SMILES notation is further converted into numerical representation for developing machine learning models using two distinct encoding representations, the MACCS (Molecular ACCess System) keys [108] and PubChem substructure fingerprints [109]. The MACCS keys consist of 166 bits representing a predefined set of common chemical fragments and topological features. To have a higher-resolution representation, we also employed the 881-bit PubChem fingerprints, which encode a broader range of hierarchic element counts and complex substructure patterns.

#### Assumptions

As part of the model development, we have assumed that Kp,uu in rats (or rodents) is representative of Kp,uu in primates (monkeys and humans). We have also assumed there is no active transport or efflux pumps involved in drug penetration to the brain.

#### Model Development

We have trained and developed both regression and classification models for predicting Kpuu. The model takes compound’s encoding representation as an input to predict the value (in regression) or label (in classification) for Kpuu. To select the best choices for featurization of compounds (drug encodings – MACCS Keys and Pubchem substructure fingerprints), machine learning algorithms (Random Forests, Extra Trees, Histogram-based Gradient Boosting), and hyperparameters for machine learning algorithm, we performed extensive parameter optimization. The information of these different parameters for drug encodings and machine learning algorithms is shown in Supplementary Table 1. The list of different hyperparameters that were optimized along with the best set of parameters is shown in Supplementary Table 2. Five-fold cross validation was utilized for optimization. Negative mean absolute error and Accuracy was used as metric for selecting parameters for regression and classification models respectively. All the hyperparameter tuning is done with random_state set as 42.

### Drug-Target Interaction (DTI) model development

#### Dataset

We utilized BindingDB [110], a publicly available extensive database providing information on drug-protein interactions, curated from various resources such as research articles, patents, etc. The dataset provides information on SMILES notation and Pubchem Compound ID (CID) for drug, and Amino Acid sequence and Uniprot ID for protein [111]. Further, the dataset quantifies drug-protein interactions using four different binding affinity metrics: IC50, Kd, Ki, and EC50. Not every drug-protein pair will have information on all binding affinity metrics.

#### Binding Affinity Metrics

The BindingDB dataset quantifies drug-protein binding by providing information on four binding affinity metrics: IC50, Kd, Ki, and EC50. IC50 (Half Maximal Inhibitory Concentration) indicates concentration of drug molecule required to inhibit biological process by 50%. Kd (Dissociation Constant) refers to the concentration of drug molecule, when 50% of the protein (or receptors) are bound. Similarly, Ki (Inhibition Constant) refers to the concentration of drug molecule or inhibitor that occupy 50% of the binding sites. EC50 (Half Maximal Effective Concentration) is a concentration of drug molecule that produces a 50 %biological response i.e. halfway between baseline and maximum possible effect.

#### Preprocessing

BindingDB dataset has information for about 3.2M drug-protein interactions. As mentioned before, not every drug-protein pair has the information on all four binding affinity metrics. Every drug-protein interaction also doesn’t provide Pubchem CID and Uniprot ID. Moreover, some drug-protein pairs have multiple entries for the same binding affinity metric. Therefore, it is apparent to perform data preprocessing steps to ensure the high quality and standard of dataset in model training and development. The data preprocessing was performed in series of three steps, as mentioned below.

#### First step

We filter out the interactions which have multiple protein chains in target sequence, along with those providing no information on Pubchem CID and Uniprot ID. It reduced the number of pairs to around 1.98M.

#### Second step

We filter out those drug-protein pairs which has different SMILES notation or Amino Acid sequences but same Pubchem CID or Uniprot ID respectively in the dataset. It led to remaining 1.27M drug-protein pairs.

### Third step

The final step in the data preprocessing, ensures that each drug-protein pair has a single entry for a particular binding affinity metric, i.e., a drug-protein pair associated with multiple values for any binding affinity metric (e.g. Kd) is removed from the dataset.

Following the data preprocessing steps, we have 270,957 pairs with Ki values, 122,861 pairs with EC50 values, 683,726 pairs with IC50 values and 32,100 pairs with Kd values. We developed machine learning models (both regression and classification) to predict all four binding affinity metrics. Further, these binding affinity metrics are determined experimentally and can vary from as low as equal to 1e-3, to as high as 1e9 in nM units. As per scientific convention, the lower value of binding affinity metric indicates higher affinity. Therefore, instead of using raw values, we utilized negative log-transformed values, i.e. we developed models to predict pKd, pKi, pIC50 and pEC50.

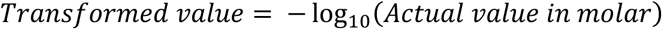

The transformation is similar to the data processing method in DeepPurpose model [34]. With this transformation, higher the transformed value, higher is the binding affinity, and vice versa. Further, binding affinity metric <= 1000 nM (transformed value >= 6) is assumed as appreciable threshold to train classification models.

#### Featurization

As mentioned above, drugs are represented by SMILES notation, and proteins are represented by Amino Acid sequence. These SMILES notation and Amino Acid sequences are converted to numerical representation via several encoding methods. Like Kpuu model development, we utilized MACCS Keys (166-dimensional) and Pubchem substructure fingerprints (881-dimensional) to encode SMILES notation into numerical representation. For the proteins, we utilized Conjoint Triad [112], Quasi Sequence order features [113], and PseudoAAC [114] to encode Amino Acid sequences into numerical representation. Conjoint Triad (343-dimensional) captures local patterns by grouping amino acids into 7 different classes based on physiochemical properties, Quasi Sequence order (100-dimensional) features capture global order i.e. long-range correlations in physiochemical properties of amino acids, and Pseudo Amino Acid Composition (30-dimensional) accounts for both the global amino acid composition and the long-range correlation of biochemical properties, such as hydrophobicity and hydrophilicity between residues.

For Conjoint Triad features, amino acids are grouped into 7 different groups (called as triads) based on their characteristics and properties followed by computing frequency of occurrence of these triads. As there are 7 different groups, the creation of triads with replacement leads to 343 features (7×7×7). The 7 groups and their corresponding amino acids are listed in Supplementary Table 4. For example, Alanine (A), Glycine (G) and Valine (V) are part of group 1, therefore, 1-1-1 (or AGV-AGV-AGV) refers to frequency of presence of either of these amino acids as three consecutive residues in the sequence.

Quasi Sequence order features account for global order, via two different distance metrics, Schneider-Wrede physicochemical distance [115] and Grantham chemical distance matrix [116]. Both distance metrics contribute 50 features each to the list of final totals 100 features. The first 20 features capture the frequence of occurrence of 20 amino acids. The other 30 features capture distances between amino acids at specific intervals varying from 1 to 30 residues, based on physiochemical properties.

Pseudo Amino Acid Composition (PseudoAAC) capture the global frequency of occurrence of different amino acids along with global long-range correlation between amino acid residues at intervals varying from 1 to 10, based on properties such as hydrophobicity, hydrophilicity, and residue mass. It leads to total of 30 features, with first 20 capturing individual amino acid frequency and remaining 10 accounting for global correlation between residues.

#### Model Development

We have trained and developed both regression and classification models for predicting all four binding affinity metrics. To select the best choices for featurization of drugs (MACCS Keys and Pubchem substructure fingerprints), proteins (Conjoint Triad, Quasi Sequence order, and PseudoAAC), machine learning algorithms (Random Forests, Extra Trees, Histogram-based Gradient Boosting), and hyperparameters for machine learning algorithm, we performed extensive parameter optimization. The information about drug and protein encodings methods along with machine learning algorithms is shown in Supplementary Table 1.

We separately tuned these parameters for classification and regression models. The parameter optimization was conducted in series of three steps. As we are dealing with a huge DTI dataset, we utilize different amount of data in each step, with it being increasing at each step. In first step, we utilize a low proportion of data (or small subset) to tune some model parameters, while in second step, we utilize a significantly higher proportion of data (3 times more data than in the first step) to further fine-tune these parameters. In third and final step, whole dataset is utilized, to select best encodings representation for both drug and protein. The information on proportion of data being used at different steps is shown in Supplementary Table 5. Due to the lowest possible feature size, we utilized the combination of MACCS Keys (for drugs) and PseudoAAC (for proteins) to tune parameters in first and second step.

As training of regression model can take significantly more time and computational resources than classification model, we initially tuned parameters for classification models and utilize those parameters as initial set of parameters (or seed) to further fine-tune regression model parameters.

#### Classification Model Parameter Optimization

The first step in parameter optimization involved tuning for parameters other than number of estimators in machine learning algorithms. The default value for number of estimators from scikit-learn [117] was utilized to train models with different parameters, followed by being optimized for their accuracy. The top five set of parameters for each machine learning algorithm were selected and used in tuning number of estimators (or iterations in case of Histogram-based Gradient Boosting) as part of second step of parameter optimization. As we tune over four different values of estimators, the second step provides us with 60 different set of parameters (3 ML models x 4 values of estimators x 5 top parameters). In third and final step, the best set of parameters from the second step is utilized to train model over the whole dataset, using different encoding representations for drug and proteins, to find the best drug and protein encoding representation for predicting a particular binding affinity metric. The list of different hyperparameters that were optimized in the series of three steps is shown in Supplementary Table 6.

#### Regression Model Parameter Optimization

The optimized set of parameters for classification model is utilized as a seed input to optimize parameters for regression. The top five set of parameters from second step of classification parameter optimization are used to optimize for the loss function (squared error, and absolute error) in first step. The top five set of parameters from first step, are used to train model in the second step using significantly higher proportion of data. The top performing set of parameters from the second step is utilized to train over the whole data, using different encoding representations for drug and protein, in step 3.

Therefore, at the end of parameter optimization, we have list of optimal parameters of both classification and regression models for predicting all four binding affinity metrics. The list of optimal parameters is shown in Supplementary Table 7. All the hyperparameter tuning is done with random_state set as 42.

### Mechanistic model for simulation of disease biomarkers

#### Systems Pharmacology Model

This study utilizes the systems pharmacology model proposed by researchers at Leiden and Merck. The model aims to analyze the changes in the amyloid precursor protein (APP) pathway after administration of BACE1 inhibitors in cisterna magna ported rhesus monkeys. The concentration of drug is retrieved from the PK model and is used to simulate the impact on the concentrations of APP metabolites Aβ42, Aβ40, soluble β-amyloid precursor protein (sAPP) α, and sAPPβ in Brain and CSF. The model accounts for the production, elimination, and brain-to-CSF transport for these metabolites. The systems-level parameters include baseline levels of these metabolites and rate constants, which we assumed and used the same values as of the research article. The other two parameters which govern the behavior and simulation of biomarker levels are defined as follows,

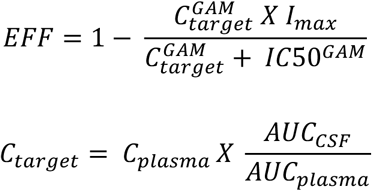

where, GAM is Hill coefficient and is assumed to be same as the research article. I_max_ i.e. maximum inhibition is assumed to be 1, while IC50 value is predicted score from the machine learning prediction (Table 3). C_plasma_ is obtained from the PK model. Further, we have assumed that in steady state, the ratio AUC_CSF_/AUC_plasma_, can be represented by Kp,uu which can be predicted using machine learning model (Table 3). Therefore, the use of IC50 and Kp,uu predictions can help us in determining EFF and C_target_ parameter values of each compound hit. The systems pharmacology model contained expressions to describe the production, elimination, and brain-to-CSF transport for the APP metabolites.

#### PK Model

The pharmacokinetics (PK) model is used to estimate the plasma concentration in the systems pharmacology model. We utilized in-house machine learning-based PK prediction framework developed by researchers at Sanofi [107] to simulate the concentration of compound hits. The model predicts the PK in rat, which is then converted to non-human primate (rhesus monkeys for systems pharmacology model) using allometric scaling [118]. Further, the dose of the compound is assumed to be 1mg/kg.

## Acknowledgements

The authors would like to thank our colleagues, Majid Vakilynejad, Qingping Wang, Frank Rapaport, Tejaswi Iyyanki, Abhinav Gupta, David Kombo, Yi Li and Madeleine Coimbra for fruitful discussions and thoughtful suggestions.

## Supplementary Figures

**Supplementary Figure 1.**
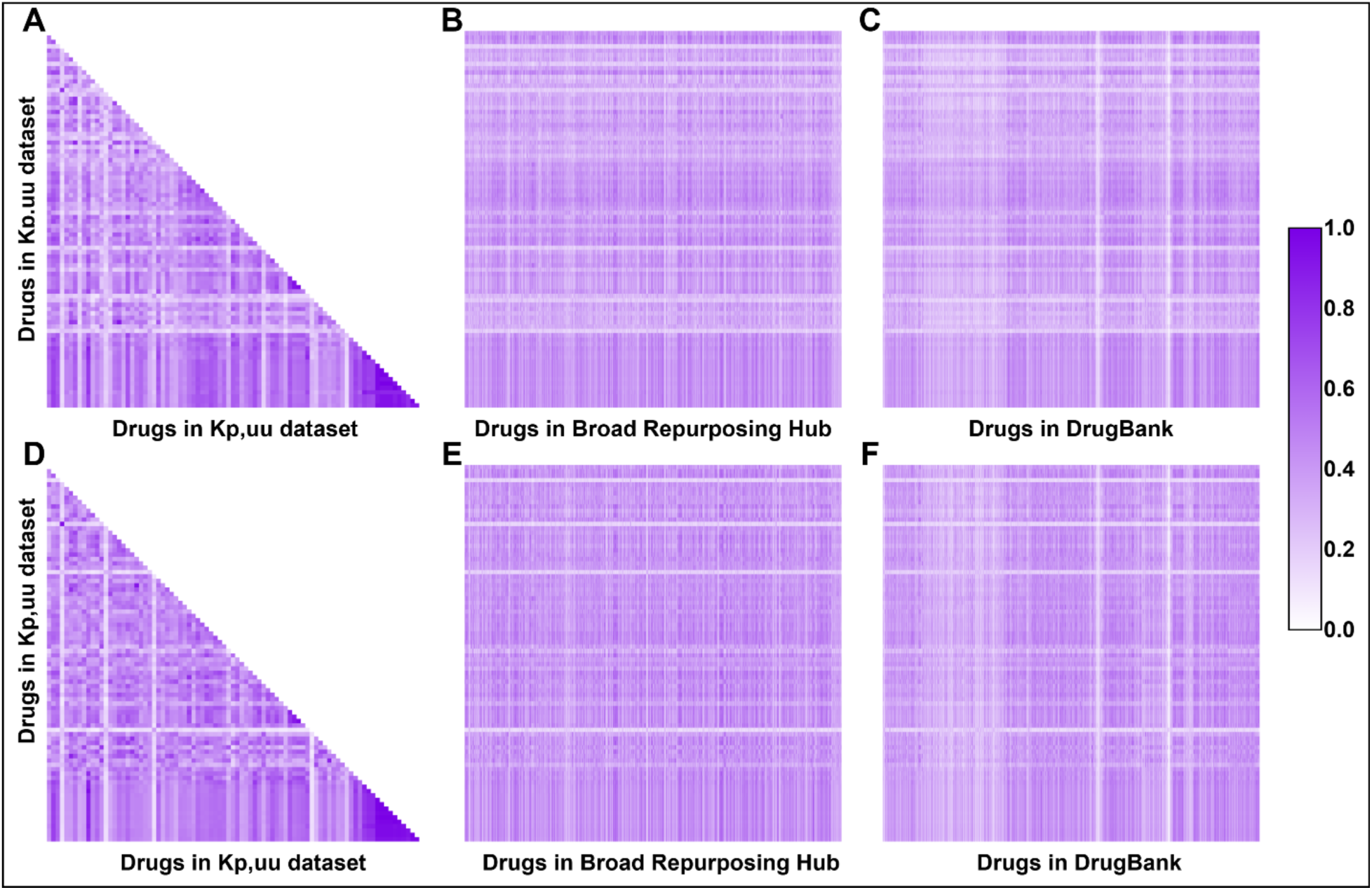
Structural similarity heatmaps. Heatmaps representing structural similarity scores of drugs in Kp,uu dataset (N = 86) among themselves, with drugs in Broad Repurposing Hub and DrugBank libraries. Heatmaps in (A)-(C) are generated using MACCS keys and (D)-(F) are generated using Pubchem substructure fingerprints.

**Supplementary Figure 2.**
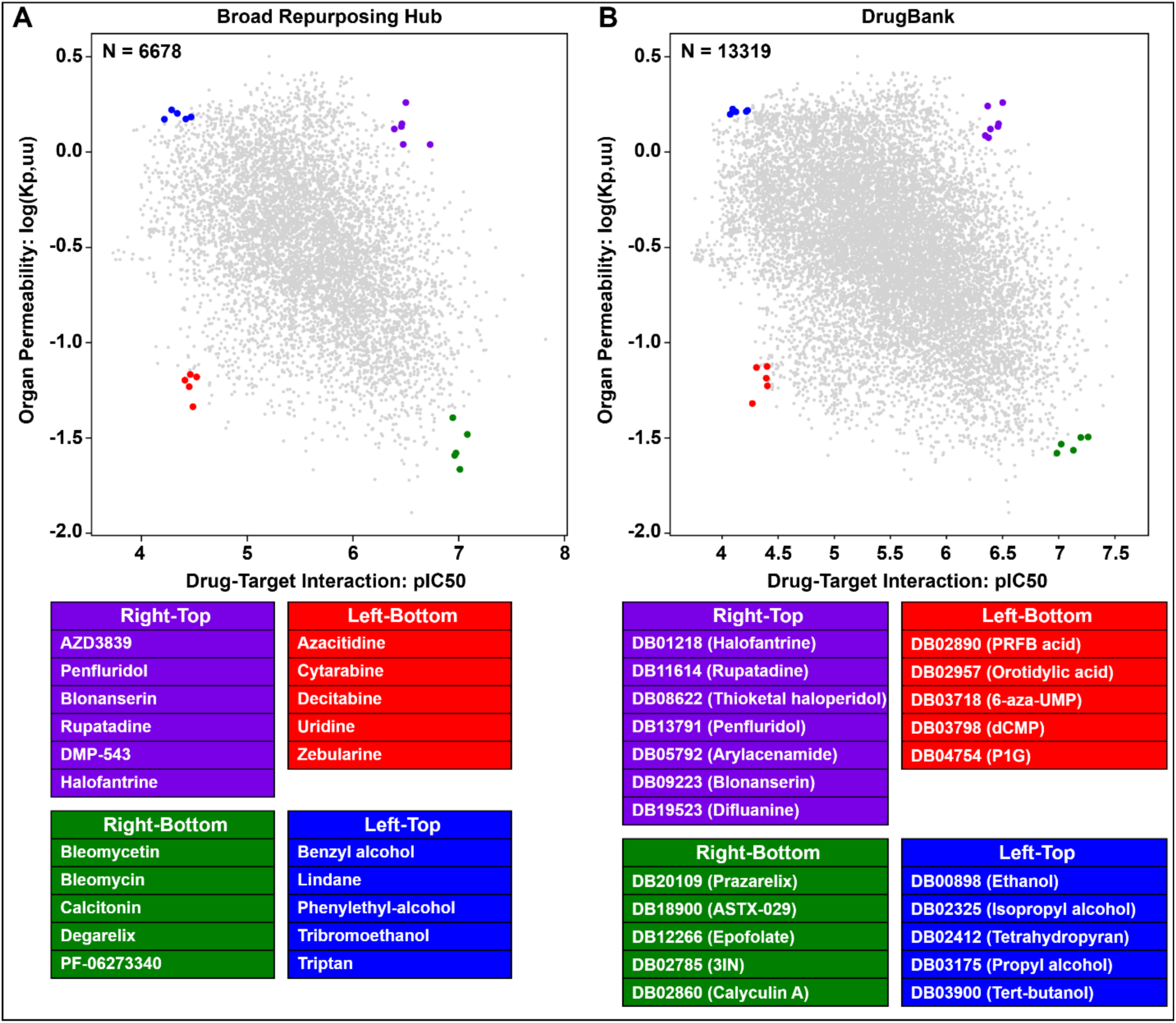
Kp,uu-DTI Landscapes highlighting selected compounds from different regions. Five compounds selected at random from Left-Bottom region (Low Kp,uu and pIC50), Right-Bottom region (Low Kp,uu and High pIC50), and Left-Top region (High Kp,uu and low pIC50) for mechanistic validation from **(A)** Broad Repurposing Hub library and **(B)** DrugBank library.

**Supplementary Figure 3.**
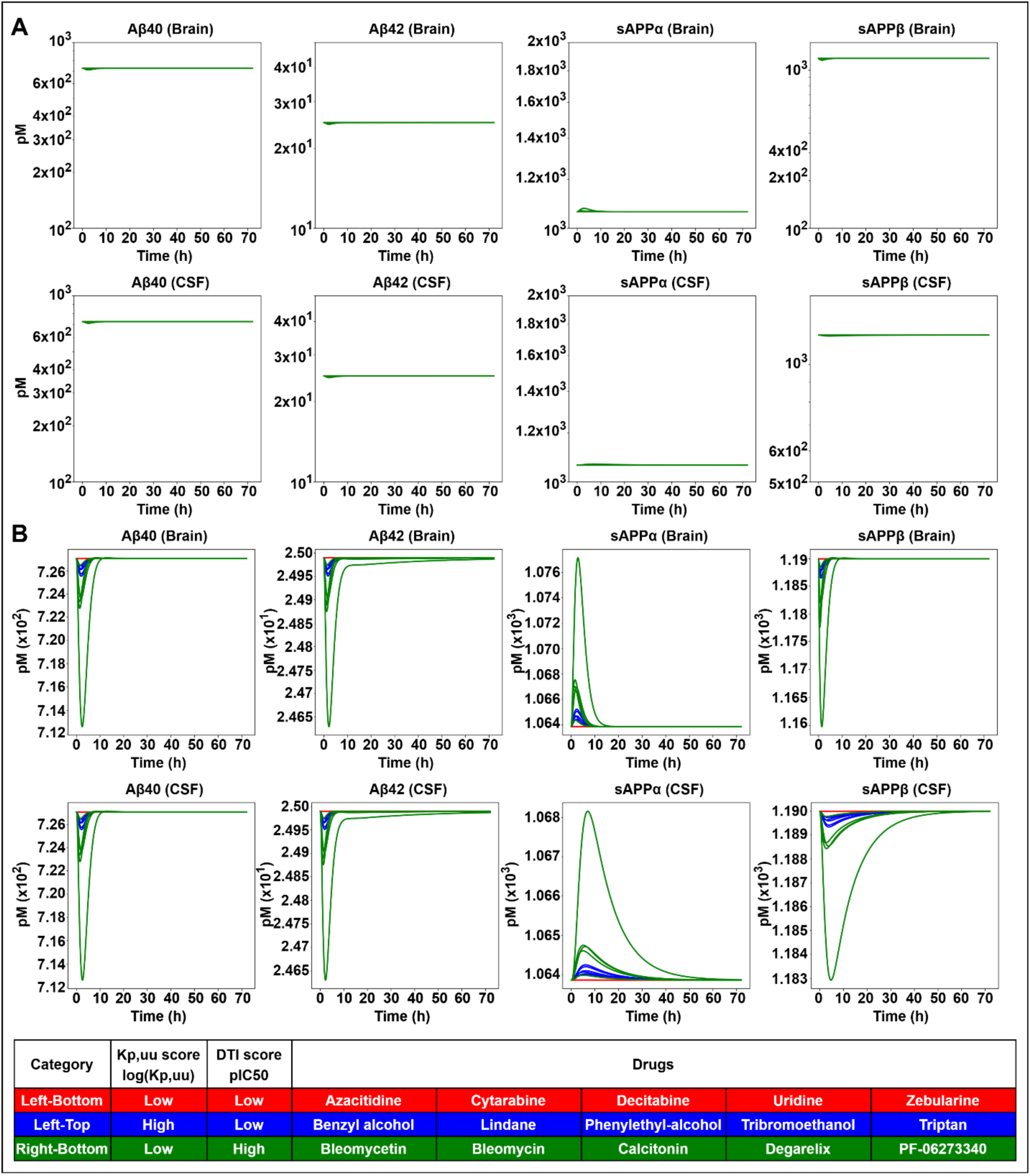
Validation of mechanistic model simulation in downstream biomarkers using non-hit compounds in Broad Repurposing Hub library. **(A)** The effect of each compound non-hit from three different regions (Left-Bottom, Left-Top and Right-Bottom) on the modulation of all four different biomarkers, in both brain and CSF region. Y-axis in all the plots have units of pM (picomolar). The scale of the y-axis is same as used in validation for compound hits (Figure 5B). **(B)** Variable scale act as magnification to understand and visualize the actual impact on downstream biomarker levels by compound non-hits.

**Supplementary Figure 4.**
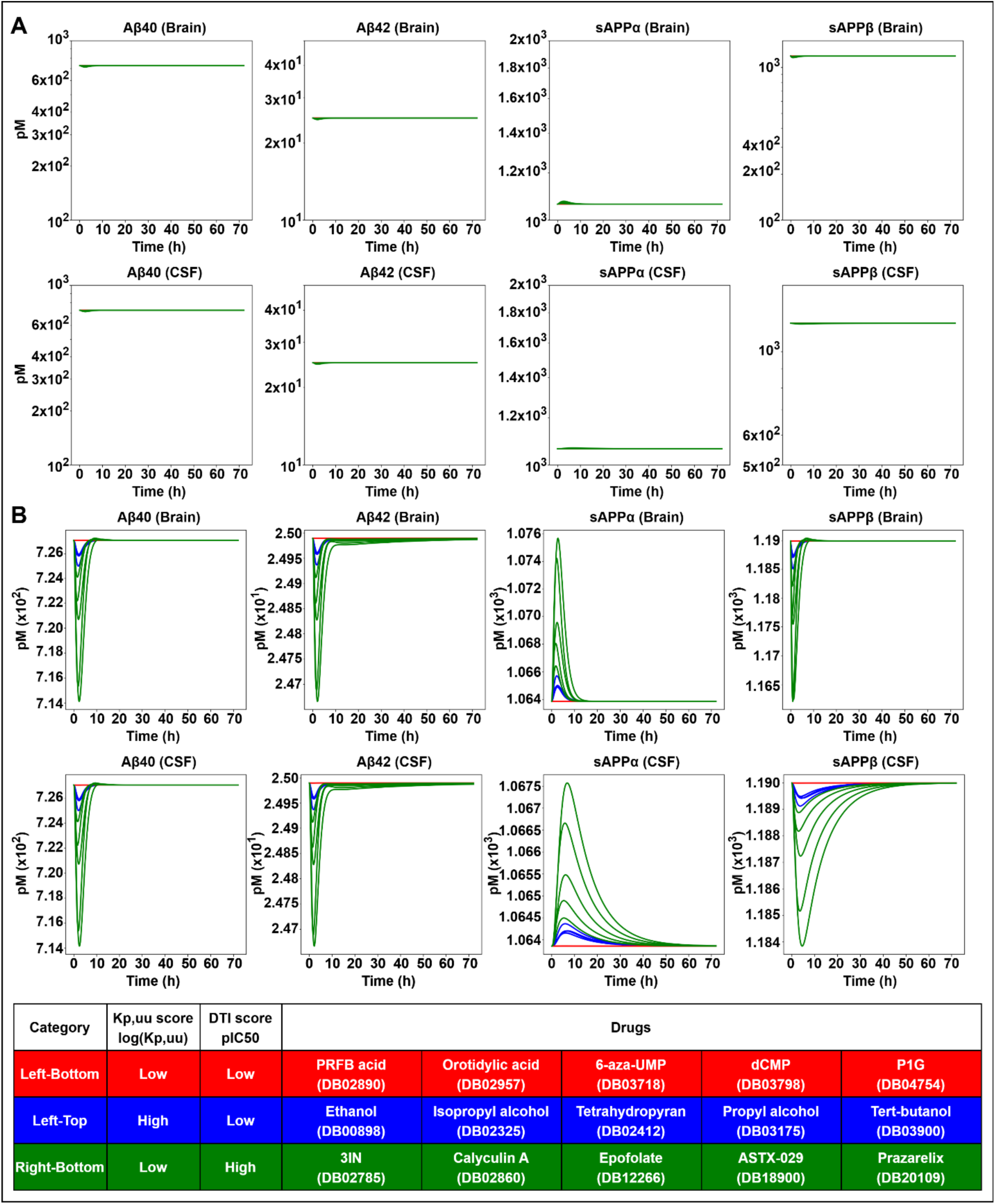
Validation of mechanistic model simulation in downstream biomarkers using non-hit compounds in DrugBank library. **(A)** The effect of each compound non-hit from three different regions (Left-Bottom, Left-Top and Right-Bottom) on the modulation of all four different biomarkers, in both brain and CSF region. Y-axis in all the plots have units of pM (picomolar). The scale of the y-axis is same as used in validation for compound hits (Figure 5B). **(B)** Variable scale act as magnification to understand and visualize the actual impact on downstream biomarker levels by compound non-hits.

## References

[1] “Research & Development.” [Online]. Available: https://www.phrma.org/policy-issues/research-development. [Accessed: 08-Feb-2026].

[2] D. Sun, W. Gao, H. Hu, and S. Zhou, “Why 90% of clinical drug development fails and how to improve it?,” Acta Pharm. Sin. B., vol. 12, no. 7, pp. 3049–3062, July 2022.

[3] P. Zhan, B. Yu, and L. Ouyang, “Drug repurposing: An effective strategy to accelerate contemporary drug discovery,” Drug Discov. Today, vol. 27, no. 7, pp. 1785–1788, July 2022.

[4] C. Gil and A. Martinez, “Is drug repurposing really the future of drug discovery or is new innovation truly the way forward?,” Expert Opin. Drug Discov., vol. 16, no. 8, pp. 829–831, Aug. 2021.

[5] V. S. Kulkarni, V. Alagarsamy, V. R. Solomon, P. A. Jose, and S. Murugesan, “Drug repurposing: An effective tool in modern drug discovery,” Russ. J. Bioorganic Chem., vol. 49, no. 2, pp. 157–166, Feb. 2023.

[6] G. Jin and S. T. C. Wong, “Toward better drug repositioning: prioritizing and integrating existing methods into efficient pipelines,” Drug Discov. Today, vol. 19, no. 5, pp. 637–644, May 2014.

[7] N. Turner, X.-Y. Zeng, B. Osborne, S. Rogers, and J.-M. Ye, “Repurposing drugs to target the diabetes epidemic,” Trends Pharmacol. Sci., vol. 37, no. 5, pp. 379–389, May 2016.

[8] S. Pushpakom et al., “Drug repurposing: progress, challenges and recommendations,” Nat. Rev. Drug Discov., vol. 18, no. 1, pp. 41–58, Jan. 2019.

[9] O.-S. Kwon, W. Kim, H.-J. Cha, and H. Lee, “In silico drug repositioning: from large-scale transcriptome data to therapeutics,” Arch. Pharm. Res., vol. 42, no. 10, pp. 879–889, Oct. 2019.

[10] A. S. Brown and C. J. Patel, “MeSHDD: Literature-based drug-drug similarity for drug repositioning,” J. Am. Med. Inform. Assoc., vol. 24, no. 3, pp. 614–618, May 2017.

[11] A. Gottlieb, G. Y. Stein, E. Ruppin, and R. Sharan, “PREDICT: a method for inferring novel drug indications with application to personalized medicine,” Mol. Syst. Biol., vol. 7, no. 1, p. 496, June 2011.

[12] J. Lamb et al., “The Connectivity Map: using gene-expression signatures to connect small molecules, genes, and disease,” Science, vol. 313, no. 5795, pp. 1929–1935, Sept. 2006.

[13] S. Sadeghi, J. Lu, and A. Ngom, “A network-based drug repurposing method via non-negative matrix factorization,” Bioinformatics, vol. 38, no. 5, pp. 1369–1377, Feb. 2022.

[14] F. Cheng et al., “Network-based approach to prediction and population-based validation of in silico drug repurposing,” Nat. Commun., vol. 9, no. 1, p. 2691, July 2018.

[15] S. Rodriguez et al., “Machine learning identifies candidates for drug repurposing in Alzheimer’s disease,” Nat. Commun., vol. 12, no. 1, p. 1033, Feb. 2021.

[16] Y. Wang et al., “DrugRepo: a novel approach to repurposing drugs based on chemical and genomic features,” Sci. Rep., vol. 12, no. 1, p. 21116, Dec. 2022.

[17] F. Ahmed et al., “SperoPredictor: An integrated machine learning and molecular docking-based drug repurposing framework with use case of COVID-19,” Front. Public Health, vol. 10, p. 902123, June 2022.

[18] H. Jia and G. C. Sosso, “Transparent machine learning model to understand drug permeability through the blood-brain barrier,” J. Chem. Inf. Model., vol. 64, no. 23, pp. 8718–8728, Dec. 2024.

[19] E. T. C. Huang et al., “Predicting blood–brain barrier permeability of molecules with a large language model and machine learning,” Sci Rep, vol. 14, no. 1, p. 15844, July 2024.

[20] S. G. Summerfield, J. W. T. Yates, and D. A. Fairman, “Free Drug Theory - no longer just a hypothesis?,” Pharm. Res., vol. 39, no. 2, pp. 213–222, Feb. 2022.

[21] E. C. M. de Lange and M. Hammarlund Udenaes, “Understanding the blood-brain barrier and beyond: Challenges and opportunities for novel CNS therapeutics,” Clin. Pharmacol. Ther., vol. 111, no. 4, pp. 758–773, Apr. 2022.

[22] I. Loryan et al., “Unbound brain-to-plasma partition coefficient, KP,uu,brain-a game changing parameter for CNS drug discovery and development,” Pharm. Res., vol. 39, no. 7, pp. 1321–1341, July 2022.

[23] A. Gupta, P. Chatelain, R. Massingham, E. N. Jonsson, and M. Hammarlund-Udenaes, “Brain distribution of cetirizine enantiomers: comparison of three different tissue-to-plasma partition coefficients: K(p), K(p,u), and K(p,uu),” Drug Metab. Dispos., vol. 34, no. 2, pp. 318–323, Feb. 2006.

[24] J. F. Morales, M. E. Ruiz, R. E. Stratford, and A. Talevi, “Application of machine learning to predict unbound drug bioavailability in the brain,” Front. Drug Discov. (Lausanne), vol. 4, no. 1360732, p. 1360732, Apr. 2024.

[25] S. Liu and Y. Kosugi, “Human brain penetration prediction using scaling approach from animal machine learning models,” AAPS J., vol. 25, no. 5, p. 86, Sept. 2023.

[26] M. Lawrenz et al., “A computational physics-based approach to predict unbound brain-to-plasma partition coefficient, KP,uu,” J. Chem. Inf. Model., vol. 63, no. 12, pp. 3786–3798, June 2023.

[27] C. Cervellati, G. Valacchi, and G. Zuliani, “BACE1 role in Alzheimer’s disease and other dementias: from the theory to the practice,” Neural Regen. Res., vol. 16, no. 12, pp. 2407–2408, Dec. 2021.

[28] H. Hampel et al., “The β-secretase BACE1 in Alzheimer’s disease,” Biol. Psychiatry, vol. 89, no. 8, pp. 745–756, Apr. 2021.

[29] R. Yan and R. Vassar, “Targeting the β secretase BACE1 for Alzheimer’s disease therapy,” Lancet Neurol., vol. 13, no. 3, pp. 319–329, Mar. 2014.

[30] G. Evin and C. Hince, “BACE1 as a therapeutic target in Alzheimer’s disease: rationale and current status,” Drugs Aging, vol. 30, no. 10, pp. 755–764, Oct. 2013.

[31] X.-Y. Meng, H.-X. Zhang, M. Mezei, and M. Cui, “Molecular docking: a powerful approach for structure-based drug discovery,” Curr. Comput. Aided Drug Des., vol. 7, no. 2, pp. 146–157, June 2011.

[32] K. Crampon, A. Giorkallos, M. Deldossi, S. Baud, and L. A. Steffenel, “Machine-learning methods for ligand-protein molecular docking,” Drug Discov. Today, vol. 27, no. 1, pp. 151–164, Jan. 2022.

[33] W. Shi, H. Yang, L. Xie, X.-X. Yin, and Y. Zhang, “A review of machine learning-based methods for predicting drug-target interactions,” Health Inf. Sci. Syst., vol. 12, no. 1, p. 30, Dec. 2024.

[34] K. Huang, T. Fu, L. M. Glass, M. Zitnik, C. Xiao, and J. Sun, “DeepPurpose: a deep learning library for drug-target interaction prediction,” Bioinformatics, vol. 36, no. 22–23, pp. 5545–5547, Apr. 2021.

[35] S. Passaro et al., “Boltz-2: Towards accurate and efficient binding affinity prediction,” bioRxiv, p. 2025.06.14.659707, 18-June-2025.

[36] K. Madrasi et al., “Systematic in silico analysis of clinically tested drugs for reducing amyloid-beta plaque accumulation in Alzheimer’s disease,” Alzheimers. Dement., vol. 17, no. 9, pp. 1487–1498, Sept. 2021.

[37] Y. Cao et al., “Neuro-dynamic quantitative systems pharmacology (qsp) model supports continued lecanemab treatment with maintenance dosing for Alzheimer’s disease,” Alzheimers. Dement., vol. 20, no. S6, p. e092093, Dec. 2024.

[38] L. Lin et al., “Quantitative systems pharmacology model for Alzheimer’s disease to predict the effect of aducanumab on brain amyloid,” CPT Pharmacometrics Syst. Pharmacol., vol. 11, no. 3, pp. 362–372, Mar. 2022.

[39] E. M. T. van Maanen et al., “Systems pharmacology analysis of the amyloid cascade after β-secretase inhibition enables the identification of an Aβ42 oligomer pool,” J. Pharmacol. Exp. Ther., vol. 357, no. 1, pp. 205–216, Apr. 2016.

[40] R. Rust, H. Yin, B. Achón Buil, A. P. Sagare, and K. Kisler, “The blood-brain barrier: a help and a hindrance,” Brain, vol. 148, no. 7, pp. 2262–2282, July 2025.

[41] J. T. Henderson and M. Piquette-Miller, “Blood-brain barrier: an impediment to neuropharmaceuticals,” Clin. Pharmacol. Ther., vol. 97, no. 4, pp. 308–313, Apr. 2015.

[42] Y. Jiao et al., “Drug delivery across the blood-brain barrier: A new strategy for the treatment of neurological diseases,” Pharmaceutics, vol. 16, no. 12, p. 1611, Dec. 2024.

[43] D. Wu, Q. Chen, X. Chen, F. Han, Z. Chen, and Y. Wang, “The blood-brain barrier: structure, regulation, and drug delivery,” Signal Transduct. Target. Ther., vol. 8, no. 1, p. 217, May 2023.

[44] A. Talevi and C. L. Bellera, “Brain-to-plasma concentration ratio and unbound partition coefficient,” in The ADME Encyclopedia, Cham: Springer International Publishing, 2021, pp. 1–6.

[45] N. Colclough, K. Chen, P. Johnström, M. Fridén, and D. F. McGinnity, “Building on the success of osimertinib: achieving CNS exposure in oncology drug discovery,” Drug Discov. Today, vol. 24, no. 5, pp. 1067–1073, May 2019.

[46] M. Gupta, J. Feng, and G. Bhisetti, “Experimental and computational methods to assess central nervous system penetration of small molecules,” Molecules, vol. 29, no. 6, p. 1264, Mar. 2024.

[47] S. J. Hindle et al., “Evolutionarily conserved roles for blood-brain barrier xenobiotic transporters in endogenous steroid partitioning and behavior,” Cell Rep., vol. 21, no. 5, pp. 1304–1316, Oct. 2017.

[48] F. Mayer, N. Mayer, L. Chinn, R. L. Pinsonneault, D. Kroetz, and R. J. Bainton, “Evolutionary conservation of vertebrate blood-brain barrier chemoprotective mechanisms in Drosophila,” J. Neurosci., vol. 29, no. 11, pp. 3538–3550, Mar. 2009.

[49] N. M. O’Brown, S. J. Pfau, and C. Gu, “Bridging barriers: a comparative look at the blood-brain barrier across organisms,” Genes Dev., vol. 32, no. 7–8, pp. 466–478, Apr. 2018.

[50] S. Lundberg and S.-I. Lee, “A unified approach to interpreting model predictions,” arXiv [cs.AI], 22-May-2017.

[51] A. Altmann, L. Toloşi, O. Sander, and T. Lengauer, “Permutation importance: a corrected feature importance measure,” Bioinformatics, vol. 26, no. 10, pp. 1340–1347, May 2010.

[52] J. Kelder, P. D. Grootenhuis, D. M. Bayada, L. P. Delbressine, and J. P. Ploemen, “Polar molecular surface as a dominating determinant for oral absorption and brain penetration of drugs,” Pharm. Res., vol. 16, no. 10, pp. 1514–1519, Oct. 1999.

[53] Q. Zhu et al., “Entropy and polarity control the partition and transportation of drug-like molecules in biological membrane,” Sci. Rep., vol. 7, no. 1, p. 17749, Dec. 2017.

[54] A. L. Hopkins and C. R. Groom, “The druggable genome,” Nat. Rev. Drug Discov., vol. 1, no. 9, pp. 727–730, Sept. 2002.

[55] M. M. Reuter et al., “Ultra-high-throughput screening of antimicrobial combination therapies using a two-stage transparent machine learning model,” bioRxivorg, p. 2024.11.25.625231, 25-Nov-2024.

[56] Y. Wang et al., “Effects of lipophilicity on the affinity and nonspecific binding of iodinated benzothiazole derivatives,” J. Mol. Neurosci., vol. 20, no. 3, pp. 255–260, Aug. 2003.

[57] M. A. Parker, D. M. Kurrasch, and D. E. Nichols, “The role of lipophilicity in determining binding affinity and functional activity for 5-HT2A receptor ligands,” Bioorg. Med. Chem., vol. 16, no. 8, pp. 4661–4669, Apr. 2008.

[58] R. Patil, S. Das, A. Stanley, L. Yadav, A. Sudhakar, and A. K. Varma, “Optimized hydrophobic interactions and hydrogen bonding at the target-ligand interface leads the pathways of drug-designing,” PLoS One, vol. 5, no. 8, p. e12029, Aug. 2010.

[59] A. R. Alhankawi et al., “The relationship between hydrophobicity and drug-protein binding in human serum albumin: A quartz crystal microbalance study,” Biophysica, vol. 2, no. 2, pp. 113–120, May 2022.

[60] J. Pollock et al., “Rational design of orthogonal multipolar interactions with fluorine in protein-ligand complexes,” J. Med. Chem., vol. 58, no. 18, pp. 7465–7474, Sept. 2015.

[61] L. Hunter, “The C-F bond as a conformational tool in organic and biological chemistry,” Beilstein J. Org. Chem., vol. 6, no. 1, p. 38, Apr. 2010.

[62] P. Bhattarai, T. A. Trombley, and R. A. Altman, “Metabolic stability of fluorinated small molecules: A physical organic chemistry perspective,” J. Med. Chem., Jan. 2026.

[63] D. Chen, N. Oezguen, P. Urvil, C. Ferguson, S. M. Dann, and T. C. Savidge, “Regulation of protein-ligand binding affinity by hydrogen bond pairing,” Sci. Adv., vol. 2, no. 3, p. e1501240, Mar. 2016.

[64] S. Y. Ablordeppey, J. B. Fischer, and R. A. Glennon, “Is a nitrogen atom an important pharmacophoric element in sigma ligand binding?,” Bioorg. Med. Chem., vol. 8, no. 8, pp. 2105–2111, Aug. 2000.

[65] L. D. Pennington and D. T. Moustakas, “The necessary nitrogen atom: A versatile high-impact design element for multiparameter optimization,” J. Med. Chem., vol. 60, no. 9, pp. 3552–3579, May 2017.

[66] L. D. Pennington, P. N. Collier, and E. Comer, “Harnessing the necessary nitrogen atom in chemical biology and drug discovery,” Med. Chem. Res., vol. 32, no. 7, pp. 1278–1293, July 2023.

[67] N. Kerru, L. Gummidi, S. Maddila, K. K. Gangu, and S. B. Jonnalagadda, “A review on recent advances in nitrogen-containing molecules and their biological applications,” Molecules, vol. 25, no. 8, p. 1909, Apr. 2020.

[68] W. Luo et al., “Nitrogen-containing heterocyclic drug products approved by the FDA in 2023: Synthesis and biological activity,” Eur. J. Med. Chem., vol. 279, no. 116838, p. 116838, Dec. 2024.

[69] A. Irfan, S. A. Al-Hussain, T. Khalid, I. Nasim, M. Mojzych, and M. E. A. Zaki, “Recent advances in clinically approved nitrogenous heterocycle-based drugs and EGFR Tyrosine kinase inhibitors for precision oncology (2020–2024): a review,” J. Saudi Chem. Soc., vol. 29, no. 6, p. 39, Dec. 2025.

[70] P. Wipf, E. M. Skoda, and A. Mann, “Conformational restriction and steric hindrance in medicinal chemistry,” in *The Practice of Medicinal Chemistry*, Elsevier, 2015, pp. 279–299.

[71] W. S. Hlavacek, R. G. Posner, and A. S. Perelson, “Steric effects on multivalent ligand-receptor binding: exclusion of ligand sites by bound cell surface receptors,” Biophys. J., vol. 76, no. 6, pp. 3031–3043, June 1999.

[72] S. M. Corsello et al., “The Drug Repurposing Hub: a next-generation drug library and information resource,” Nat. Med., vol. 23, no. 4, pp. 405–408, Apr. 2017.

[73] C. Knox et al., “DrugBank 6.0: The DrugBank knowledgebase for 2024,” Nucleic Acids Res., vol. 52, no. D1, pp. D1265–D1275, Jan. 2024.

[74] “Multibrand.” [Online]. Available: https://openinnovation.astrazeneca.com/home/preclinical-research/preclinical-molecules/azd3839.html. [Accessed: 09-Mar-2026].

[75] “Donepezil.” [Online]. Available: https://go.drugbank.com/drugs/DB00843. [Accessed: 07-Feb-2026].

[76] B. Seltzer, “Donepezil: a review,” Expert Opin. Drug Metab. Toxicol., vol. 1, no. 3, pp. 527–536, Oct. 2005.

[77] Q. Huang, C. Liao, F. Ge, J. Ao, and T. Liu, “Acetylcholine bidirectionally regulates learning and memory,” J. Neurorestoratology, vol. 10, no. 2, p. 100002, June 2022.

[78] J. Haam and J. L. Yakel, “Cholinergic modulation of the hippocampal region and memory function,” J. Neurochem., vol. 142 Suppl 2, no. S2, pp. 111–121, Aug. 2017.

[79] Z.-R. Chen, J.-B. Huang, S.-L. Yang, and F.-F. Hong, “Role of cholinergic signaling in Alzheimer’s disease,” Molecules, vol. 27, no. 6, p. 1816, Mar. 2022.

[80] O. Delbono, Z.-M. Wang, and M. L. Messi, “The rise and deceleration of neuronal excitability in aging and Alzheimer’s disease: Mechanisms, implications, and therapeutic targets,” Ageing Res. Rev., vol. 114, no. 102999, p. 102999, Jan. 2026.

[81] H. Targa Dias Anastacio, N. Matosin, and L. Ooi, “Neuronal hyperexcitability in Alzheimer’s disease: what are the drivers behind this aberrant phenotype?,” Transl. Psychiatry, vol. 12, no. 1, p. 257, June 2022.

[82] T. J. Baumgartner et al., “Inhibition of the GSK3β/Nav1.6 complex suppresses early-stage Alzheimer’s hyperexcitability,” Alzheimers. Dement., vol. 21, no. 7, p. e70507, July 2025.

[83] Y. Wang, Y. Shi, and H. Wei, “Calcium dysregulation in Alzheimer’s disease: A target for new drug development,” J. Alzheimers Dis. Parkinsonism, vol. 7, no. 5, pp. 1–7, Aug. 2017.

[84] A. Kumar, K. Bodhinathan, and T. C. Foster, “Susceptibility to Calcium Dysregulation during Brain Aging,” Front. Aging Neurosci., vol. 1, p. 2, Nov. 2009.

[85] N. C. Wildburger, A. Lin-Ye, M. A. Baird, D. Lei, and J. Bao, “Neuroprotective effects of blockers for T-type calcium channels,” Mol. Neurodegener., vol. 4, no. 1, p. 44, Oct. 2009.

[86] F. G. Sanchez-Conde, E. N. Jimenez-Vazquez, D. S. Auerbach, and D. K. Jones, “The ERG1 K+ channel and its role in neuronal health and disease,” Front. Mol. Neurosci., vol. 15, p. 890368, May 2022.

[87] Open Resources for Nursing (Open RN), K. Ernstmeyer, and E. Christman, “Chapter 6 Psychotropic Medications,” in Nursing: Mental Health and Community Concepts [Internet], Chippewa Valley Technical College, 2022.

[88] M. D. Wood and P. B. Wren, “Serotonin-dopamine interactions: implications for the design of novel therapeutic agents for psychiatric disorders,” Prog. Brain Res., vol. 172, pp. 213–230, Jan. 2008.

[89] A. V. McCormick, J. M. Wheeler, C. R. Guthrie, N. F. Liachko, and B. C. Kraemer, “Dopamine D2 receptor antagonism suppresses tau aggregation and neurotoxicity,” Biol. Psychiatry, vol. 73, no. 5, pp. 464–471, Mar. 2013.

[90] M. J. Ramírez, “5-HT6 receptors and Alzheimer’s disease,” Alzheimers. Res. Ther., vol. 5, no. 2, p. 15, Apr. 2013.

[91] M. Andrews, B. Tousi, and M. N. Sabbagh, “5HT6 antagonists in the treatment of Alzheimer’s dementia: Current progress,” Neurol. Ther., vol. 7, no. 1, pp. 51–58, June 2018.

[92] S. Hashimoto and T. C. Saido, “Critical review: involvement of endoplasmic reticulum stress in the aetiology of Alzheimer’s disease,” Open Biol., vol. 8, no. 4, p. 180024, Apr. 2018.

[93] B. D. Roussel, A. J. Kruppa, E. Miranda, D. C. Crowther, D. A. Lomas, and S. J. Marciniak, “Endoplasmic reticulum dysfunction in neurological disease,” Lancet Neurol., vol. 12, no. 1, pp. 105–118, Jan. 2013.

[94] A. Ajoolabady, D. Lindholm, J. Ren, and D. Pratico, “ER stress and UPR in Alzheimer’s disease: mechanisms, pathogenesis, treatments,” Cell Death Dis., vol. 13, no. 8, p. 706, Aug. 2022.

[95] T.-Y. Weng, S.-Y. A. Tsai, and T.-P. Su, “Roles of sigma-1 receptors on mitochondrial functions relevant to neurodegenerative diseases,” J. Biomed. Sci., vol. 24, no. 1, p. 74, Sept. 2017.

[96] T.-P. Su, T. Hayashi, T. Maurice, S. Buch, and A. E. Ruoho, “The sigma-1 receptor chaperone as an inter-organelle signaling modulator,” Trends Pharmacol. Sci., vol. 31, no. 12, pp. 557–566, Dec. 2010.

[97] W. Zhang, D. Xiao, Q. Mao, and H. Xia, “Role of neuroinflammation in neurodegeneration development,” Signal Transduct. Target. Ther., vol. 8, no. 1, p. 267, July 2023.

[98] A. Adamu, S. Li, F. Gao, and G. Xue, “The role of neuroinflammation in neurodegenerative diseases: current understanding and future therapeutic targets,” Front. Aging Neurosci., vol. 16, p. 1347987, Apr. 2024.

[99] A. E. Musto and M. Samii, “Platelet-activating factor receptor antagonism targets neuroinflammation in experimental epilepsy: PAF-r Antagonism Targets Neuroinflammation,” Epilepsia, vol. 52, no. 3, pp. 551–561, Mar. 2011.

[100] Z. Mputhia, E. Hone, T. Tripathi, T. Sargeant, R. Martins, and P. Bharadwaj, “Autophagy modulation as a treatment of amyloid diseases,” Molecules, vol. 24, no. 18, p. 3372, Sept. 2019.

[101] Y. Fu et al., “Activating autophagy to eliminate toxic protein aggregates with small molecules in neurodegenerative diseases,” Pharmacol. Rev., vol. 77, no. 3, p. 100053, May 2025.

[102] R. Schwaha, “The similarity principle – new trends and applications in ligand-based drug discovery and ADMET profiling,” Sci. Pharm., vol. 76, no. 1, pp. 5–18, Mar. 2008.

[103] D. Bajusz, A. Rácz, and K. Héberger, “Why is Tanimoto index an appropriate choice for fingerprint-based similarity calculations?,” J. Cheminform., vol. 7, no. 1, p. 20, May 2015.

[104] E. Heuer, R. F. Rosen, A. Cintron, and L. C. Walker, “Nonhuman primate models of Alzheimer-like cerebral proteopathy,” Curr. Pharm. Des., vol. 18, no. 8, pp. 1159–1169, 2012.

[105] R. F. Rosen et al., “Comparative pathobiology of β-amyloid and the unique susceptibility of humans to Alzheimer’s disease,” Neurobiol. Aging, vol. 44, pp. 185–196, Aug. 2016.

[106] M. B. Podlisny, D. R. Tolan, and D. J. Selkoe, “Homology of the amyloid beta protein precursor in monkey and human supports a primate model for beta amyloidosis in Alzheimer’s disease,” Am. J. Pathol., vol. 138, no. 6, pp. 1423–1435, June 1991.

[107] N. Pillai, A. Abos, D. Teutonico, and P. D. Mavroudis, “Machine learning framework to predict pharmacokinetic profile of small molecule drugs based on chemical structure,” Clin. Transl. Sci., vol. 17, no. 5, p. e13824, May 2024.

[108] J. L. Durant, B. A. Leland, D. R. Henry, and J. G. Nourse, “Reoptimization of MDL keys for use in drug discovery,” J. Chem. Inf. Comput. Sci., vol. 42, no. 6, pp. 1273–1280, Nov. 2002.

[109] S. Kim et al., “PubChem 2023 update,” Nucleic Acids Res., vol. 51, no. D1, pp. D1373–D1380, Jan. 2023.

[110] T. Liu et al., “BindingDB in 2024: a FAIR knowledgebase of protein-small molecule binding data,” Nucleic Acids Res., vol. 53, no. D1, pp. D1633–D1644, Jan. 2025.

[111] UniProt Consortium, “UniProt: The universal protein knowledgebase in 2023,” Nucleic Acids Res., vol. 51, no. D1, pp. D523–D531, Jan. 2023.

[112] J. Shen et al., “Predicting protein-protein interactions based only on sequences information,” Proc. Natl. Acad. Sci. U. S. A., vol. 104, no. 11, pp. 4337–4341, Mar. 2007.

[113] K. C. Chou, “Prediction of protein subcellular locations by incorporating quasi-sequence-order effect,” Biochem. Biophys. Res. Commun., vol. 278, no. 2, pp. 477–483, Nov. 2000.

[114] K. C. Chou, “Prediction of protein cellular attributes using pseudo-amino acid composition,” Proteins, vol. 43, no. 3, pp. 246–255, May 2001.

[115] G. Schneider and P. Wrede, “The rational design of amino acid sequences by artificial neural networks and simulated molecular evolution: de novo design of an idealized leader peptidase cleavage site,” Biophys. J., vol. 66, no. 2 Pt 1, pp. 335–344, Feb. 1994.

[116] R. Grantham, “Amino acid difference formula to help explain protein evolution,” Science, vol. 185, no. 4154, pp. 862–864, Sept. 1974.

[117] F. Pedregosa et al., “Scikit-learn: Machine Learning in Python,” arXiv [cs.LG], no. 85, pp. 2825–2830, 02-Jan-2012.

[118] Z. Lu, R. Kaspera, Y. Naritomi, and T. Wang, “Dose finding in single dose studies by allometric scaling,” in Drug Discovery and Evaluation: Methods in Clinical Pharmacology, Cham: Springer International Publishing, 2018, pp. 1–11.

