## Supplementary tables for "A Modular Machine Learning Framework for Small-Molecule Drug Repurposing Based on Organ Permeability, Target Binding, and Biomarker Modulation"

**Supplementary Table 1. Choices utilized for Drug encodings, Protein encodings, and Machine learning algorithms for hyperparameter optimization**

| | |
|---|---|
| Drug Encodings | MACCS Keys |
| | Pubchem substructure fingerprints |
| Protein Encodings | PseudoAAC |
| | Conjoint Triad |
| | Quasi Sequence Order |
| Machine Learning Algorithms | Random Forests |
| | Extra Trees |
| | Histogram-based Gradient Boosting |

| Supplementary Table 2. List of hyperparameters and their corresponding values that were optimized for machine learning algorithms, along with the best performing model for regression task for Kp,uu prediction. | |
|---|---|
| Random Forests | n_estimators: 5, 10, 50, 100, 200, 500<br>criterion: 'squared_error', 'friedman_mse', 'absolute_error'<br>max_depth: 3, 5, 7, 9, None<br>min_samples_split: 2, 5, 10<br>min_samples_leaf: 1, 2, 4<br>max_features: 'sqrt', 'log2', 0.25, 0.5, 0.75, 1.0<br>bootstrap: True, False |
| Extra Trees | n_estimators: 5, 10, 50, 100, 200, 500<br>criterion: 'squared_error', 'friedman_mse', 'absolute_error'<br>max_depth: 3, 5, 7, 9, None<br>min_samples_split: 2, 5, 10<br>min_samples_leaf: 1, 2, 4<br>max_features: 'sqrt', 'log2', 0.25, 0.5, 0.75, 1.0<br>bootstrap: True, False |
| Histogram-based Gradient Boosting | max_iter: 5, 10, 50, 100, 200, 500<br>loss: 'squared_error', 'absolute_error'<br>learning_rate: 0.02, 0.05, 0.1<br>max_bins: 64, 128, 255<br>max_leaf_nodes: 32, 64, 128, 256<br>min_samples_leaf: 10, 20, 50, 100<br>l2_regularization: 0.0, 1e-4, 1e-3, 1e-2<br>max_depth: None, 8, 12, 16 |
| **Best Model** | Drug Encoding: MACCS Keys<br>Machine Learning Algorithm: Extra Trees<br>bootstrap: True<br>criterion: 'absolute_error'<br>max_depth: 7<br>max_features: 'sqrt'<br>min_samples_leaf: 1<br>min_samples_split: 2<br>n_estimators: 10 |

| **Supplementary Table 3. List of hyperparameters and their corresponding values that were optimized for machine learning algorithms, along with the best performing model for classification task for Kp,uu prediction.** | |
|---|---|
| Random Forests | n_estimators: 5, 10, 50, 100, 200, 500<br>criterion: 'gini', 'entropy'<br>max_depth: 3, 5, 7, 9, None<br>min_samples_split: 2, 5, 10<br>min_samples_leaf: 1, 2, 4<br>max_features: 'sqrt', 'log2', 0.25, 0.5, 0.75, 1.0<br>bootstrap: [True, False |
| Extra Trees | n_estimators: 5, 10, 50, 100, 200, 500<br>criterion: 'gini', 'entropy'<br>max_depth: 3, 5, 7, 9, None<br>min_samples_split: 2, 5, 10<br>min_samples_leaf: 1, 2, 4<br>max_features: 'sqrt', 'log2', 0.25, 0.5, 0.75, 1.0<br>bootstrap: [True, False |
| Histogram-based Gradient Boosting | max_iter: 5, 10, 50, 100, 200, 500<br>learning_rate: 0.02, 0.05, 0.1<br>max_bins: 64, 128, 255<br>max_leaf_nodes: 32, 64, 128, 256<br>min_samples_leaf: 10, 20, 50, 100<br>l2_regularization: 0.0, 1e-4, 1e-3, 1e-2<br>max_depth: None, 8, 12, 16 |
| **Best Model** | Drug Encoding: MACCS Keys<br>Machine Learning Algorithm: Extra Trees<br>bootstrap: True<br>criterion: 'gini'<br>max_depth: 9<br>max_features: 0.5<br>min_samples_leaf: 1<br>min_samples_split: 5<br>n_estimators: 10 |

**Supplementary Table 4. Categorization of amino acids into different groups based on characteristics and properties.**

| Group | Amino Acids | Characteristics |
|---|---|---|
| 1 | Alanine (A) | Small, nonpolar |
| | Glycine (G) | |
| | Valine (V) | |
| 2 | Isoleucine (I) | Large, nonpolar |
| | Leucine (L) | |
| | Phenylalanine (F) | |
| | Proline (P) | |
| 3 | Tyrosine (Y) | Polar, uncharged |
| | Methionine (M) | |
| | Threonine (T) | |
| | Serine (S) | |
| 4 | Histidine (H) | Large, polar/aromatic |
| | Asparagine (N) | |
| | Glutamine (Q) | |
| | Tryptophan (W) | |
| 5 | Arginine (R) | Positively charged |
| | Lysine (K) | |
| 6 | Aspartic Acid/Aspartate (D) | Negatively charged |
| | Glutamic Acid/Glutamate (E) | |
| 7 | Cysteine (C) | Unique sulfur bonding |

| **Supplementary Table 5. Proportion (or percentage) of data being used in first and second step of hyperparameter tuning of DTI models** | | | | |
|---|---|---|---|---|
| Step | EC50 | IC50 | Kd | Ki |
| First | 0.15 | 0.05 | 0.3 | 0.1 |
| Second | 0.45 | 0.15 | 0.39 | 0.3 |

| Supplementary Table 6. List of hyperparameters and their corresponding values that were optimized for machine learning algorithms in series of three steps for DTI prediction models. | | |
|---|---|---|
| First Step | Random Forests | criterion: 'gini', 'entropy', 'log_loss'<br>max_depth: 3, 5, 7, 9<br>min_samples_split: 2, 5, 10, 20<br>min_samples_leaf: 1, 2, 4, 8<br>max_features: 'sqrt', 'log2', 0.2, 0.5, 0.8, 1.0<br>bootstrap: True, False<br>class_weight : None, 'balanced' |
| | Extra Trees | criterion: 'gini', 'entropy', 'log_loss'<br>max_depth: 3, 5, 7, 9<br>min_samples_split: 2, 5, 10, 20<br>min_samples_leaf: 1, 2, 4, 8<br>max_features: 'sqrt', 'log2', 0.2, 0.5, 0.8, 1.0<br>bootstrap: True, False<br>class_weight: None, 'balanced' |
| | Histogram-based Gradient Boosting | learning_rate: 0.02, 0.05, 0.1<br>max_bins: 64, 128, 255<br>max_leaf_nodes: 32, 64, 128, 256<br>min_samples_leaf: 10, 20, 50, 100<br>l2_regularization: 0.0, 1e-4, 1e-3, 1e-2<br>max_depth: None, 8, 12, 16 |
| Second Step | Number of Estimators/Iterations | 50, 100, 200, 500 |
| Third Step | Drug Encodings | MACCS Keys<br>Pubchem substructure fingerprints |
| | Protein Encodings | PseudoAAC<br>Conjoint Triad<br>Quasi Sequence Order |

| **Supplementary Table 7. List of hyperparameters chosen for best performing models for classification task for predicting all four DTI binding affinity metrics (pEC50, pIC50, pKi, and pKd).** | | | | |
|---|---|---|---|---|
| Binding Affinity Metric | Drug Encoding | Protein Encoding | Machine Learning algorithm | Hyperparameters |
| EC50 | Pubchem substructure fingerprints | Conjoint Triad | Histogram-based Gradient Boosting Algorithm | max_bins=64<br>max_depth=16<br>max_iter=500<br>max_leaf_nodes=256 |
| IC50 | | | | l2_regularization=0.0001<br>max_bins=64<br>max_depth=12<br>max_iter=500<br>max_leaf_nodes=256 |
| Ki | | Quasi Sequence Order | | max_bins=64<br>max_depth=16<br>max_iter=500<br>max_leaf_nodes=256<br>min_samples_leaf=50 |
| Kd | | | | l2_regularization: 0.0001<br>max_depth: 12<br>max_iter: 200<br>max_leaf_nodes: 128<br>min_samples_leaf=10 |

| Supplementary Table 8. List of hyperparameters chosen for best performing models for regression task for predicting all four DTI binding affinity metrics (pEC50, pIC50, pKi, and pKd). | | | | |
|---|---|---|---|---|
| Binding Affinity Metric | Drug Encoding | Protein Encoding | Machine Learning algorithm | Hyperparameters |
| EC50 | Pubchem substructure fingerprints | Conjoint Triad | Histogram-based Gradient Boosting Algorithm | l2_regularization=0.01<br>max_bins=128<br>max_iter=500<br>max_leaf_nodes=256 |
| IC50 | | | | l2_regularization=0.01<br>max_bins=64<br>max_depth=16<br>max_iter=500<br>max_leaf_nodes=256<br>min_samples_leaf=50 |
| Ki | | | | l2_regularization=0.0001<br>max_bins=64<br>max_depth=16<br>max_iter=500<br>max_leaf_nodes=256 |
| Kd | | Quasi Sequence Order | | l2_regularization=0.01<br>max_iter=500<br>max_leaf_nodes=128 |

**Supplementary Table 9. Performance metrics for Regression and Classification models for DTI prediction for Kd, Ki, and EC50.**

| | | Kd | Ki | EC50 |
|---|---|---|---|---|
| Regression Performance Metrics | Spearman correlation (R) | 0.83 | 0.85 | 0.9 |
| | P-value | ~0 | ~0 | ~0 |
| | Pearson correlation | 0.86 | 0.86 | 0.9 |
| | Mean Absolute Error (MAE) | 0.52 | 0.6 | 0.52 |
| | Mean Squared Error (MSE) | 0.59 | 0.64 | 0.54 |
| | Coefficient of Determination (R2) | 0.74 | 0.73 | 0.8 |
| Regression-based Classification Metrics | Accuracy | 0.87 | 0.86 | 0.88 |
| | F1-score | 0.83 | 0.9 | 0.88 |
| | Sensitivity | 0.81 | 0.93 | 0.89 |
| | Matthews Correlation Coefficient (MCC) | 0.73 | 0.68 | 0.76 |
| | Area Under Curve - ROC | 0.86 | 0.83 | 0.88 |
| Classification Performance Metrics | Accuracy | 0.87 | 0.88 | 0.88 |
| | F1-score | 0.82 | 0.91 | 0.89 |
| | Sensitivity | 0.79 | 0.93 | 0.89 |
| | Matthews Correlation Coefficient (MCC) | 0.72 | 0.71 | 0.77 |
| | Area Under Curve - ROC | 0.94 | 0.94 | 0.95 |

| Supplementary Table 10. The list of compounds hits identified by M3LDR framework, along with information on molar mass, predicted Kp,uu and IC50 scores, and computed clearance and volume of distribution from PK model, to be given as an input to the mechanistic model for downstream modulation of biomarkers. | | | | | | |
|---|---|---|---|---|---|---|
| Name | DrugBank ID | Molar Mass | Kp,uu | IC50 (uM) | CL (ml/min) | V (ml) |
| AZD3839 | - | 431.422 | 1.09 | 0.19 | 15.09 | 1,144.26 |
| Blonanserin | DB09223 | 367.512 | 1.36 | 0.35 | 18.09 | 3,515.38 |
| DMP-543 | - | 412.4 | 1.1 | 0.33 | 10.64 | 1,101.83 |
| Halofantrine | DB01218 | 500.424 | 1.32 | 0.41 | 15.87 | 1,719.59 |
| Penfluridol | DB13791 | 523.97 | 1.41 | 0.34 | 8.79 | 2,267.25 |
| Rupatadine | DB11614 | 415.97 | 1.82 | 0.32 | 20.50 | 1,969.12 |
| Difluanine | DB19523 | 449.26 | 1.22 | 0.45 | 11.39 | 3,812.34 |
| Arylacenamide | DB05792 | 396.437 | 1.74 | 0.43 | 29.10 | 1,925.73 |
| Thioketal haloperidol | DB08622 | 452.048 | 1.19 | 0.42 | 17.75 | 944.83 |
